# Isolation and characterisation of Nipah virus neutralising candidate therapeutic monoclonal antibodies from an mRNA-immunised pig

**DOI:** 10.64898/2026.08.28.745669

**Authors:** Miriam Pedrera, Nichakorn Pipatpadungsin, Darwyn Kobasa, Ahmed M.E. Elrefaey, Barbara Holzer, Rebecca K. McLean, Bryce Warner, Robert Vendramelli, Nazia Thakur, Robert Stass, Jack W. P. Hayes, Lobna Medfai, Joshua E. Sealy, Sylvia Crossley, John C. Schwartz, Danish Munir, William Mwangi, Dalan Bailey, Thang Truong, Elma Tchilian, Bradley Pickering, Thomas A. Bowden, Simon P. Graham

**Author notes:** The authors contributed equally to this work (Miriam Pedrera, Nichakorn Pipatpadungsin, Darwyn Kobasa). Centro de Investigación en Sanidad Animal, Instituto de Investigación y Tecnología Agraria y Alimentaria, Consejo Superior de Investigaciones Científicas (CISA-INIA-CSIC), Valdeolmos, Madrid, Spain.

## Abstract

Nipah virus (NiV) is a highly pathogenic zoonotic paramyxovirus with epidemic potential. Despite the threat NiV poses, no therapeutics are licensed to treat infection. Studies have shown that monoclonal antibodies (mAb) can protect animals against NiV and the related Hendra virus (HeV). The best studied mAb, m102.4, has been used to treat infected patients on a compassionate basis, and has entered clinical trials. However, there is a need to define additional mAbs with therapeutic potential, which could be combined with m102.4 to improve neutralising potency and breadth. Here, we isolated five high affinity mAbs from a pig vaccinated with mRNA encoding the NiV G glycoprotein, which bound the recombinant NiV Malaysia strain (NiV-M) G, and one of which (mAb A2) also cross-reacted with HeV G. All mAbs neutralised NiV-M pseudovirus but only mAb A2 neutralised pseudovirus representing the NiV Bangladesh (NiV-B) strain. mAb A2 and the most potent NiV-M neutralising mAb, C1, showed minimal competition with each other and m102.4, suggesting recognition of non-overlapping epitopes. Single-particle cryogenic electron microscopy of the NiV-M G receptor binding domain complexed to A1 and C2 Fab fragments revealed distinct epitopes that did not overlap with the receptor-binding site, targeted by m102.4, suggesting action through steric impedance of receptor binding or interference downstream of receptor engagement. Administration of mAb A2 to hamsters did not provide complete protection against NiV-B challenge (60% survival), however, a split dose of mAb A2 and m102.4 provided the same protection as m102.4 alone (100% survival). Collectively, these data demonstrate the potential of the porcine model for isolation of therapeutic candidate mAbs, which contribute both to our understanding of the NiV G antigenic landscape, and the development of mAb combinations, that exert complementary mechanisms of neutralisation, for therapeutic intervention.

## Introduction

Nipah virus (NiV; *Henipavirus nipahense*) is a highly pathogenic member of the family *Paramyxoviridae*. NiV poses a significant zoonotic threat because of its broad host range and widespread distribution of *Pteropus spp.* bats, which serve as a natural reservoir. NiV infection causes a severe respiratory and neurologic disease in humans, with case-fatality rates ranging from 40% to over 75% [1]. NiV is one of the most concerning threats to public health security in South and Southeast Asia, driven by continued population growth and increasing human interactions with NiV-infected wildlife as urban development expands into natural ecosystems. Humans may become infected by contact with infected bat secretions, with subsequent human-to-human spread. Human infections also occur following exposure to infected pigs, where NiV causes a less severe respiratory and neurologic syndrome [2,3]. As demonstrated by the Ebola outbreaks in Africa, there is potential for highly pathogenic emerging infections that normally induce small isolated and contained outbreaks, to cause large epidemics with significant mortality and morbidity. The number of people at risk from NiV infection is estimated to be over two billion. Despite the importance of NiV as an emerging disease with pandemic potential, no vaccines nor therapeutics are approved. Consequently, the WHO lists Nipah virus infection as a priority for research and development in public health emergency contexts.

Experimental studies have shown that treatment with monoclonal antibodies (mAbs) can protect animals against lethal infections with NiV and the related Hendra virus (HeV). In this context, the best studied mAb, m102.4 [4–6], directed against the receptor-binding domain (RBD) of the henipaviral attachment glycoprotein (G), has previously been used to treat infected humans on a compassionate basis, and has been evaluated in a successful phase I clinical trial [7]. As demonstrated for other viral infections e.g., Ebola virus [8], there is a benefit to using mAb ‘cocktails’ to improve both the potency and breadth of neutralising activity. There is therefore a need to define additional NiV mAbs with therapeutic potential, which could be combined with m102.4 [7]. To support this goal, recent studies have reported the isolation of neutralising humanised mAbs targeting the NiV fusion (F) glycoprotein, an alternative therapeutic target, which can provide protection in animal models [9–11]. Furthermore, a recent study isolated F and G-specific mAbs from humanised mice, which neutralised both NiV and HeV, and showed that a combination of two mAbs provided increased resistance against a suite of escape mutant pseudoviruses [12].

Animal models that can mimic human disease are essential for developing henipavirus vaccines and therapeutics [13]. Among the existing animal models, the Syrian golden hamster is the most widely used small animal model for NiV infection, which develops clinical symptoms and pathologic features that closely resemble disease in humans [14]. However, large animal models, such as pigs, have been shown to more accurately predict vaccine outcome in humans [15]. NiV vaccine candidates have been evaluated for immunogenicity and efficacy in pigs [16–20]. In addition to providing preclinical data to support human vaccine development [21], these studies have been conducted with a view to developing a vaccine for pigs as a primary amplifying host of NiV [20]. Immunised or infected pigs can also be used to isolate candidate therapeutic mAbs. Influenza A virus-neutralising haemagglutinin-specific mAbs were isolated from infected pigs that targeted epitopes similar to those recognised by human antibodies [22,23]. These studies highlight the utility of the pig as a source of relevant antibodies, as well as a valuable model for the evaluation and delivery of therapeutic mAbs [24,25].

mAbs recognising different epitopes may exert distinct functional effects and act synergistically to prevent or control virus infection [26,27]. Furthermore, targeting multiple epitopes may provide broader protection against viral variants and reduce the risk of viral escape mutations [27,28]. Isolation of NiV-specific mAbs from pigs may not only provide valuable insights into protective epitopes but also provide promising therapeutic candidates that can be humanised for clinical applications. We report here the isolation of mAbs from a pig immunised with mRNA expressing NiV G, the characterisation of two neutralising epitopes, and the evaluation of protective efficacy of the lead candidate mAb in comparison to, and in combination with, mAb m102.4.

## Materials and methods

### Ethics statement

A vaccine study using pigs was carried out at the Animal and Plant Health Agency, Addlestone, UK, in accordance with UK Animals (Scientific Procedures) Act 1986 (Project License P9C86DC55) and with approval from the local Animal Welfare and Ethical Review Body (AWERB; approval number P986D55-1-002). A mAb efficacy study using Syrian golden hamsters was performed at the National Microbiology Laboratory (NML) of the Public Health Agency of Canada. Experiments were approved by the Animal Care Committee at the Canadian Science Centre for Human and Animal Health per the guidelines provided by the Canadian Council on Animal Care under animal use document C-23-004, and with approval from The Pirbright Institute AWERB. All infectious work with NiV Bangladesh (NiV-B) strain was performed in the containment level 4 laboratory at the NML. All hamsters were monitored and weighed daily throughout the experiment and were provided food and water *ad libitum*.

### Immunisation of pigs with NiV vaccine candidates

A vaccine immunogenicity study with pigs was carried out as described previously [16,20]. Briefly, groups (n = 5 or 6) of 8-10-week-old, female, Large White-Landrace-Hampshire cross-bred pigs were immunised intramuscularly at 0 and 21 days post-vaccination (dpv) with a lipid nanoparticle formulated synthetic nucleoside-modified mRNA encoding a soluble version of the G protein from NiV Malaysia strain (GenBank AY029768) (NiV-M sG) [20] or bovine herpes virus 4 (BoHV-4) vectors expressing NiV-M G or F glycoproteins [16]. Blood samples were collected weekly from 0 ‒ 42 dpv for serum and peripheral blood mononuclear cells (PBMC) isolation. PBMC were routinely isolated from heparinised blood by density gradient centrifugation, and the cells were either used immediately in RPMI-1640 medium supplemented with 10% FBS, 100 IU/mL penicillin, and 100 µg/mL streptomycin (all Thermo Fisher Scientific) (cRPMI) for use in assays or cryopreserved in 10% DMSO (Merck Life Science) in FBS for later analysis [16].

### Indirect ELISA to detect henipavirus glycoprotein-specific antibodies

Porcine serum antibody and mAb binding to henipavirus glycoproteins were assessed by indirect ELISA as described [16]. Briefly, ELISA plates were coated with 100 ng/well recombinant soluble NiV and HeV sG proteins [18]. After blocking with 5% skimmed milk in PBS, two-fold serial dilutions of porcine serum samples or recombinant mAbs were incubated. Wells were incubated with horseradish peroxidase (HRP)–conjugated rabbit polyclonal antibody against porcine IgG (Merck), before addition of TMB solution (Merck) and the subsequent addition of 2N sulfuric acid stop solution. Absorbance was measured at 450 nm in a GloMax® Multi+ Detection plate reader (Promega).

### Porcine IgG ELISpot assay

To determine the frequency of circulating NiV G-specific antibody secreting B cells, an IgG ELISpot assay was performed on freshly isolated porcine PBMC. Ethanol pre-treated multiscreen 96-well plates (MAIPS4510; Millipore) were pre-coated with 15 µg/mL purified anti-porcine IgG mAb (Clone MT421, Mabtech) diluted in PBS and incubated overnight at 4°C. After washing with PBS, plates were blocked with cRPMI for 1 h at 37°C. Blocking solution was removed and PBMC were plated at 5 × 10^5^ cells/well in cRPMI for antigen-specific IgG detection and negative controls or at 5 × 10^4^ cells/well for total IgG detection. After 18 h incubation at 37 °C/5% CO_2_, medium was removed, and the remaining cells were lysed with cold distilled water and washed with PBS. For the total IgG control wells, biotinylated anti-porcine IgG mAb (MT424-biotin, Mabtech) was added at 0.5 µg/mL in 0.5% FBS in PBS (incubation buffer). To detect NiV G-specific IgG, recombinant protein NiV-M sG and the irrelevant keyhole limpet hemocyanin protein (KLH; Merck) was biotinylated using the EZ-link Micro Sulfo-NHS-Biotinylation Kit (Thermo Fisher Scientific). Protein biotinylation was confirmed and quantified using the Pierce Biotin Quantitation Kit (Thermo Fisher Scientific) and protein concentration determined using the Pierce BCA Protein Assay Kit (Thermo Fisher Scientific). Biotinylated NiV-M sG and KLH were added to relevant wells at a final concentration of 2.5 μg/mL in incubation buffer. Each condition was tested in triplicate. Following an incubation period of 2 h at room temperature (RT), plates were washed with PBS, and binding revealed with streptavidin–ALP enzyme conjugate (Mabtech) in incubation buffer for 1 h at RT in the dark, washing and addition of BCIP/NBTplus substrate (Mabtech). Plates were incubated at RT in the dark for 10 min or until visible spots developed. The reaction was stopped with water and, after removing the plastic cover, both sides of the plate were thoroughly washed. The numbers of spots were determined using an AID iSpot reader (Autoimmun Diagnostika GmbH, Germany) and data were expressed as the number of antibody-secreting cells (ASC) per million PBMC.

### Flow cytometric analysis of NiV sG tetramer-stained porcine B cells

To detect circulating NiV G-specific B cells after immunisation, fresh and previously cryopreserved PBMC were stained with NiV G tetramers in combination with flow cytometry. NiV G tetramers were assembled by incubating biotinylated NiV-M sG protein with streptavidin-Brilliant Violet (BV) 421 or streptavidin-BV650 (both BioLegend) at a molar ratio of 4:1. A negative control decoy tetramer using biotinylated SARS-CoV-2 spike protein RBD [29] was assembled using streptavidin-PerCP-Cy5.5 (BioLegend). Tetramers were assembled a minimum of 24 h in advance and stored at 4°C in the dark. PBMC were adjusted at 1 × 10^6^ cells/well in 96-well round bottom plates and were stained first with the decoy tetramer diluted in cold PBS. All incubation steps were for 30 min on ice in the dark. After washing twice with 1% FBS/PBS, a combination of NiV G-BV421 and NiV G-BV650 tetramers in cold PBS were added to the cells and incubated for 30 min. After washing, cells were stained with Zombie near-IR fixable viability dye (BioLegend), biotinylated mAbs against lineage markers (anti-porcine CD3ε-biotin, clone PPT3; anti-porcine CD8α-biotin, clone 76-2-11, both Southern Biotech; and anti-human CD14-biotin, clone REA599, Miltenyi Biotec), anti-porcine IgG-Alexa Fluor-647 mAb (Cohesion Biosciences, Generon), IgA-FITC polyclonal Ab (Bio-Rad Antibodies) and IgM-PE mAb (clone K52 1C3, Bio-Rad Antibodies; conjugated to PE with the Lightning-Link^®^ PE Ab Labelling Kit, Abcam). After 30 min incubation, cells were washed and stained with streptavidin-BV605 (BioLegend). Finally, after washing and a fixation step with 4% paraformaldehyde at 4°C, cells were acquired using a BD LSR Fortessa cell analyser and the data analysed using FlowJo software (BD Biosciences). Single colour controls were used for compensation and fluorescence minus one tetramer controls were used to set thresholds. Each sample was stained in triplicate wells. Samples were gated on lymphocytes (SSC-A vs FSC-A) and then singlet cells (FSC-H vs FSC-A). Following exclusion of dead cells, non-B cells (determined as CD3/CD14/CD8α^+^ cells) and decoy tetramer^+^ cells, the percentages of IgG^+^ or IgM^+^ cells labelled with both NiV G tetramers were assessed (Supplementary Figure 1).

### Single cell sorting of NiV G-binding IgG^+^ B cells

150 mL of blood was collected in Alsever’s solution (Merck) from three pigs immunised with the mRNA vectored NiV G vaccine candidate at 28 dpv (7 days post-boost). Blood was incubated overnight at RT with continuous rocking and PBMC isolated. B cells were enriched by depletion of monocytes, T cells and NK cells using the biotinylated CD14, CD3 and CD8α mAbs described above. PBMC were incubated with 10 μL of each mAb per 10^8^ cells and per mL of 2% FBS/PBS during 20 min at 4°C, washed once with 2% FBS/PBS and incubated for 20 min at 4°C with 100 μL/10^8^ cells of anti-biotin magnetic microbeads (Miltenyi Biotec) and per mL of 2% FBS, 2mM EDTA in PBS (MACS buffer). After washing once with 2% FBS/PBS, cells were resuspended in 0.5mL MACS buffer per 10^8^ cells, passed through a cell strainer (Miltenyi Biotec), and applied to pre-equilibrated LD columns (Miltenyi Biotec). The flow through, collected as the B cell enriched fraction, was centrifugated and cells resuspended in 2%FBS/PBS, cells enumerated using a MACSQuant Analyzer (Miltenyi Biotec).

Enriched B cells were stained with the NiV G-BV421 tetramer followed by the addition of the surface marker antibodies as described above, with the exception that Zombie Near-IR was replaced with Zombie Aqua fixable viability dye (BioLegend), the biotinylated lineage-specific mAbs as well as the IgA-FITC Ab were omitted, and the streptavidin-BV605 was replaced with the streptavidin-BV650 (BioLegend). After the last incubation, stained cells were washed and adjusted at 10^7^/mL in RPMI-1640 medium supplemented with 2% FBS. Stained cells were passed through a nylon mesh-top blue cap on a Falcon™ round-bottom test tube (Thermo Fisher Scientific) to ensure a single cell suspension and put on ice prior to sorting. Cells were gated on lymphocytes and, following the exclusion of dead cells and non-B cells, NiV G tetramer^+^ IgG^+^ cells were sorted individually (Supplementary Figure 2) using a FACS Aria III cell sorter (BD Biosciences) into hard shell 96-well PCR plates (Bio-Rad) containing 10 μL/well of 10mM Tris pH 7.4 with RNasin (Promega, UK) as described [22]. After sorting, plates were sealed, centrifuged for 5 min at 300 × *g*, at 4°C and then stored at −80°C.

### Cloning and expression of recombinant porcine mAbs

A two-step RT-PCR method to amplify the variable gene segments of the IgG heavy and light chains before cloning into porcine expression vectors as described [22].

Briefly, cDNA was synthesised from the single sorted cells and aliquots used in PCRs to amplify VDJ heavy (H) or VJ light (L) genes. A nested PCR method was used to amplify the H and kappa chains, whereas only one PCR was used to amplify the lambda chain. PCR products were then analysed on an agarose gel, and either gel extracted or directly purified and used as the template for the second PCR. H chain PCR products were cloned into a KpnI/PstI-linearised expression vector containing the porcine IgG1 constant domain (pNeoSec-SsFc-IgG1). The kappa and lambda L chain PCR products were cloned into linearised porcine expression vectors containing the constant domain of the respective L chain; pNeoSec-SsLC-κ and pNeoSec-SsLC-λ, respectively. Plasmid DNA was purified (QIAprep Spin Miniprep Kit, Qiagen) and Sanger sequenced. Native pairs of H and L chain expressing plasmids were subsequently transiently transfected into Expi293F cells (Thermo Fisher Scientific), culture supernatant harvested and IgG purified by affinity chromatography [22]. Purified recombinant porcine mAbs were buffer exchanged to PBS with 0.01% sodium azide (Merck) and stored at 4°C. The concentration of mAbs were determined using a porcine IgG ELISA kit (ELISA Flex: Porcine IgG (HRP); Mabtech). Recombinant porcine influenza A virus hemagglutinin-specific mAb (PB21; IgG1) [22] and human mAb m102.4 (expression plasmids kindly provided by Ariel Isaacs, Keith Chappell and Daniel Watterson, University of Queensland, Brisbane, Australia) were similarly expressed and used as controls in *in vitro* assays.

To produce larger quantities of mAbs for evaluation *in vivo,* expression plasmids were transiently co-transfected into ExpiCHO cells (Thermo Fisher Scientific). Purified mAbs were buffer exchanged by gel filtration or dialysis in PBS, filter sterilised by passing through a 0.2 µm glass-fibre/cellulose acetate syringe filter and tested for endotoxin (Pierce™ Chromogenic Endotoxin Quant Kit, Thermo Fisher Scientific).

For structural analyses, selected antibodies were expressed as Fab as described [30]. Briefly VDJ H genes were ordered as synthetic gene blocks (IDT) and cloned into a porcine IgG1 Fab expression (pNeoSec-SsVH-IgG1) vector carrying CH1 constant domain of IgG1 fused to a six histidine (6xHis) tag at the C-terminus and transiently co-expressed with respective light chains in Expi293F cells as described for full length antibodies. The resulting Fab fragments were purified by nickel affinity chromatography on the ÄKTA pure™ 25 using a 5mL HisTrap column (Cytiva), and then buffer exchanged by dialysis into PBS without calcium and magnesium. Concentration of the purified Fab was determined by A_280_ divided by extinction coefficient calculated based on the amino acid sequence of the corresponding protein and diluted to 1 mg/mL.

### Biolayer interferometry

Biolayer interferometry assays were performed on an Octet RED96 instrument using streptavidin (SA) biosensors (both Sartorius) to determine NiV-M sG-mAb binding kinetics at the sensor surface. SA biosensors were loaded into a 96-well black plate (Greiner) and pre-hydrated in HBS-EP buffer (GE Healthcare). Biotinylated goat anti-pig IgG (Fc) antibody (Bio-Rad Antibodies) was immobilised at 50 nM onto the SA biosensors. This was then used to capture the porcine mAbs at 10 nM. Finally, association and dissociation of recombinant NiV-M sG protein was determined using a concentration range of 15 - 0.11 nM. All steps were carried out in HBS-EP (GE Healthcare), with association and dissociation lasting 2000 s each step. Data were analysed using the Octet Analysis Software v12.2.1.23 (Sartorius).

### NiV pseudovirus neutralisation assay

Neutralising activity of the NiV-specific porcine mAbs was assessed using NiV-M and - B pseudovirus particles (NiV-Mpp and NiV-Bpp) as described [31]. In brief, serial dilutions of mAbs were incubated with NiV-Mpp or NiV-Bpp in 96-well, flat bottomed, white opaque plates (Thermo Fisher Scientific) at 37°C for 1 h. HEK-293T cells were added and plates incubated for a further 72 h. Luciferase activity was measured using Luciferase Assay Substrate and GloMax Multi+ Detection System (both Promega). Pseudovirus neutralisation titres were calculated as the inverse of the mAb concentration which showed a 50% inhibition of luciferase values (IC_50_), compared to no mAb controls.

### NiV cell fusion neutralisation assay

The ability of mAbs to neutralise NiV glycoprotein-mediated cell-cell fusion was assessed as described [16]. In brief, serial dilutions of mAbs were incubated with HEK293 cells, transduced lentivirus expressing half of a split Renilla luciferase-GFP reporter and transfected with plasmids expressing F and G from either NiV-M or -B, in 96-well, flat bottomed, white opaque plates (Thermo Fisher Scientific) at 37°C for 1 h. HEK293T target cells, transduced to express the other half of the split Renilla luciferase-GFP reporter, were added and incubated for a further 18 h. Luciferase activity was measured as described above. Fusion neutralising titres were calculated as the inverse of the mAb concentration which showed a 50% inhibition of luciferase values (IC_50_), compared to no mAb controls.

### Epitope binning by competitive ELISA

A competitive ELISA was performed to evaluate mAb competition for binding to NiV-M sG. ELISA plates were coated with recombinant NiV-M sG as described above. Plates were blocked with 100 µL/well of 2% bovine serum albumin in 0.1 M PBS containing 0.1% Tween 20 (blocking solution) and incubated for 2 h at 37°C. Plates were washed with PBS with 0.05% Tween 20 (PBS-T). 50 µL of non-biotinylated mAb at 10 µg/mL, diluted in 2% blocking solution, was added and plates incubated for 1 h at 37°C and then washed with PBS-T. 50 µL of mAb biotinylated using the EZ-link Micro Sulfo-NHS-Biotinylation Kit (Thermo Fisher Scientific) were added as two-fold serial dilutions starting from 1 µg/mL in blocking solution. Plates were incubated for 1 h at 37°C and then washed with PBS-T. 50 µL of streptavidin-HRP (Thermo Fisher Scientific) diluted 1:1000 in blocking solution, was added to the plates and incubated for 1 h at 37°C. Plates were washed and 100 µL of TMB substrate was added to the plates for 5 min at room temperature, after which the reaction was stopped by addition of 100 µL 0.5 N sulfuric acid and absorbance at 450 nm measured. Background signal was determined as the mean absorbance from control wells containing the antibody without the corresponding biotinylated antibody. The background-corrected absorbance values were used to calculate the percentage of residual binding of each biotinylated antibody in the presence of competing unlabelled antibodies.

### NiV G receptor binding domain production

The NiV-M G RBD (residues 183−602, GenBank NC_002728) was cloned into the pHLsec vector with a KHHHHHH C-terminal tag [32], as previously described [33,34]. NiV-M G RBD was recombinantly produced in Expi293F cells (Thermo Fisher Scientific) following transient transfection with polyethylenimine (PEI) using a DNA:PEI mass ratio of 1:5 in the presence of an α-mannosidase inhibitor, kifunensine, at a final concentration of 5 µM. Cell supernatant was diafiltrated against 10 mM Tris-Cl pH 8.0 150 mM NaCl buffer using ÄKTA Flux™ system (Cytiva), affinity purified using a HisTrap HP column (Cytiva), and deglycosylated with endoglycosylaseF_1_ (75 µg per mg protein, 16 h, 21°C). Protein was purified by size-exclusion chromatography using a Superdex 200 10/300 GL Increase column (Cytiva) equilibrated with 10 mM Tris-Cl pH 8.0, 150 mM NaCl buffer. Prior to single particle cryogenic electron microscopy (cryo-EM) sample preparation, NiV G RBD was complexed with a three-fold molar excess of both Fab A2 and Fab C1. The NiV G-Fab A2/Fab C1 complex was separated from unbound excess Fab by size-exclusion chromatography using a Superdex 200 10/300 GL increase column (Cytiva) equilibrated with 10 mM Tris-Cl pH 8.0, 150 mM NaCl buffer.

### Cryo-EM sample preparation and data collection

QuantiFoil™ R1.2/1.3 grid with 2 nm continuous carbon was glow-discharged using a Harrick Plasma Cleaner for 30 s. The NiV G-Fab A2/Fab C1 complex was applied onto the glow-discharged grid prior to immediate plunge-freezing using Vitrobot Mark IV (Thermo Fisher Scientific) with −15 blot force for 3.5 s. The grid was clipped with a Nanosoft autogrid ring and c-clip and stored at liquid nitrogen temperature until data collection. Data collection was performed using a 300 kV Krios II microscope at the electron Bio-Imaging Centre (eBIC), Diamond Light Source, Didcot, UK, equipped with Schottky X-FEG electron source and Gatan K3 24 megapixels detector (5760 x 4092). The collection was performed with a 100 µm objective aperture in position, 50.00 e^-^/Å^2^ electron dose equally fractionated over 50 frames, and 105,000 magnification (0.831 Å/pixel). 12,777 movies were collected in total.

### Cryo-EM data processing and model building

The collected movies were motion-corrected using MotionCor2 [35] wrapped in PATo autoprocessing pipeline at eBIC [36]. The motion-corrected micrographs were further processed in Cryosparc v5.0.6 [37]. Patch CTF was used to estimate the contrast transfer function (CTF) for each micrograph [37]. The particles were initially picked using a blob picker with an elliptical mask with 90 Å and 120 Å wide on the short and long axes, respectively. 7,052,049 particles were extracted at 128-pixel box size and subjected to 2D classifications. After iterative 2D classifications, 170,323 particles from good classes were selected as templates for another round of particle picking. Using the template picker, 13,955,283 particles were extracted at 256-pixel box size which was then cropped to 64-pixel box size in Fourier space. After several rounds of 2D classifications, 975,932 particles from good classes were extracted at 384-pixel box size. An additional round of 2D classification was performed and the number of good particles reduced to 578,289. *Ab-initio* reconstruction with 2 classes was performed to obtain an initial set of 3D maps. The best map was chosen for non-uniform refinement [38], and local refinement with a mask focussing on the complex density before being subjected to heterogeneous refinement along with other maps using the whole particle set. This pipeline was then performed iteratively until the best map resolution and features stopped improving. The final map was resolved at 3.19 Å (FSC=0.143) resolution with 119,065 particles. A summary of cryo-EM data collection and refinement statistics are presented in Supplementary Table 1 and data processing in Supplementary Figure 3. An initial model was built with ModelAngelo [39], and the model was refined iteratively with COOT v0.9 [40] and Phenix.RealspaceRefine [41]. The model geometry was validated using Molprobity webserver [42].

### Evaluation of the protective efficacy of mAbs in vivo

A study was conducted to evaluate the prophylactic effectiveness of the porcine mAb A2, either alone or in combination with mAb m102.4, to protect hamsters from disease caused by NiV-B infection. Twenty-four, mixed sex, Syrian golden hamsters, 4-6 weeks of age (60-100 g body weight), were randomly assigned to four treatment groups (n=6, with 3 males and 3 females per group) and acclimatised for 7 days prior to the start of the experiment. All procedures were performed with isoflurane inhalation anaesthesia. On −1 days post-infection (dpi), hamsters were inoculated by intraperitoneal injection with PBS (group 1), 1 mg mAb m102.4 in PBS (group 2), 1 mg mAb A2 in PBS (group 3), and 0.5 mg of mAb m102.4 and 0.5 mg Ab A2 in PBS (group 4). All animals were challenged on 0 dpi via intranasal inoculation with NiV-B at a dose of 10^5^ TCID_50_ in a volume of 0.1 mL. Animals were monitored for clinical signs and scored to the severity of disease and euthanised at humane endpoints [43] or upon experiment completion (19 dpi). On 3 and 5 dpi, oral, nasal, and rectal swab samples were collected. Swabs were pre-wetted in DMEM, placed in cryovials containing 1 mL DMEM after sampling, and frozen at −80 °C until analysis.

### Virus and titrations

Recombinant NiV-B was generated using reverse genetics as described [43]. VeroE6 cells (ATCC) and were cultured in DMEM supplemented with 5% bovine growth serum (HyClone™ BGS; Cytiva) and 100 IU/mL penicillin, and 100 µg/mL (Thermo Fisher Scientific) for maintenance. Virus titration was performed on VeroE6 cells seeded in 96 well tissue culture plates in DMEM; 1% BGS; 100 IU/mL penicillin, and 100 µg/mL (viral infection medium). Incubation conditions for VeroE6 cells were 37°C with 5% CO_2_. Infectious NiV-B titres in stock virus and oral, nasal, and rectal swabs were determined by TCID_50_ assay. Briefly, samples were thawed, vortexed, and spun at 1500 × *g* for 10 min. A ten-fold serial dilution of the samples was made in viral infection medium, and dilutions were added to 96-well plates of 95-100% confluent VeroE6 cells in triplicate. The cells were incubated at 37 °C with 5% CO_2_ for 5 days, and cytopathic effect was scored and titres determined as TCID_50_ [44].

### Data analysis

GraphPad Prism 11 (GraphPad Software) was used for graphical and statistical analysis of data sets. Flow cytometry data was analysed using FlowJo software (BD Biosciences). Statistical differences were analysed using a one- or two-way ANOVA or a mixed-effects model followed by a Sidak’s or Tukey’s multiple comparison test to compare treatment groups, timepoints or mAbs. *p*-values < 0.05 were considered statistically significant.

## Results

### Monitoring NiV glycoprotein-specific plasma B cells following immunisation

NiV-M G-specific plasma B cell responses in pigs vaccinated with mRNA and BoHV-4 vectors were evaluated longitudinally by IgG ELISpot assay (Figure 1A, B). NiV G-specific IgG-secreting cells were observed in both NiV G vaccinated groups (mRNA-NiV G and BoHV-4 NiV G), while absent in the BoHV-4 NiV F group (Figure 1A). In contrast, IgG responses to the irrelevant antigen (KLH) were minimal in all three vaccine groups at all timepoints tested (Figure 1B). In both NiV G vaccinated groups, there was an increase in the number of NiV G-specific IgG antibody-secreting cells (ASC) from 7 dpv, although this only achieved statistical significance compared to the BoHV-4 NiV F group on certain timepoints i.e., BoHV-4 NiV G group only on 7 dpv whereas the mRNA NiV G group on 21, 28, and 42 dpv. Whilst not statistically significant due to inter-animal variation, there was evidence of a plasmablast burst at 28 dpv/7 days post-boost, which was more pronounced in the mRNA NiV G group. Flow cytometric assessment of B cell binding to NiV-M sG tetramers at 28 dpv/7 days post-boost revealed elevated frequencies of labelled IgM^+^ and IgG^+^ B cells within PBMC from the mRNA G vaccinated group, albeit without statistical significance (Figure 1C)

**Figure 1.**
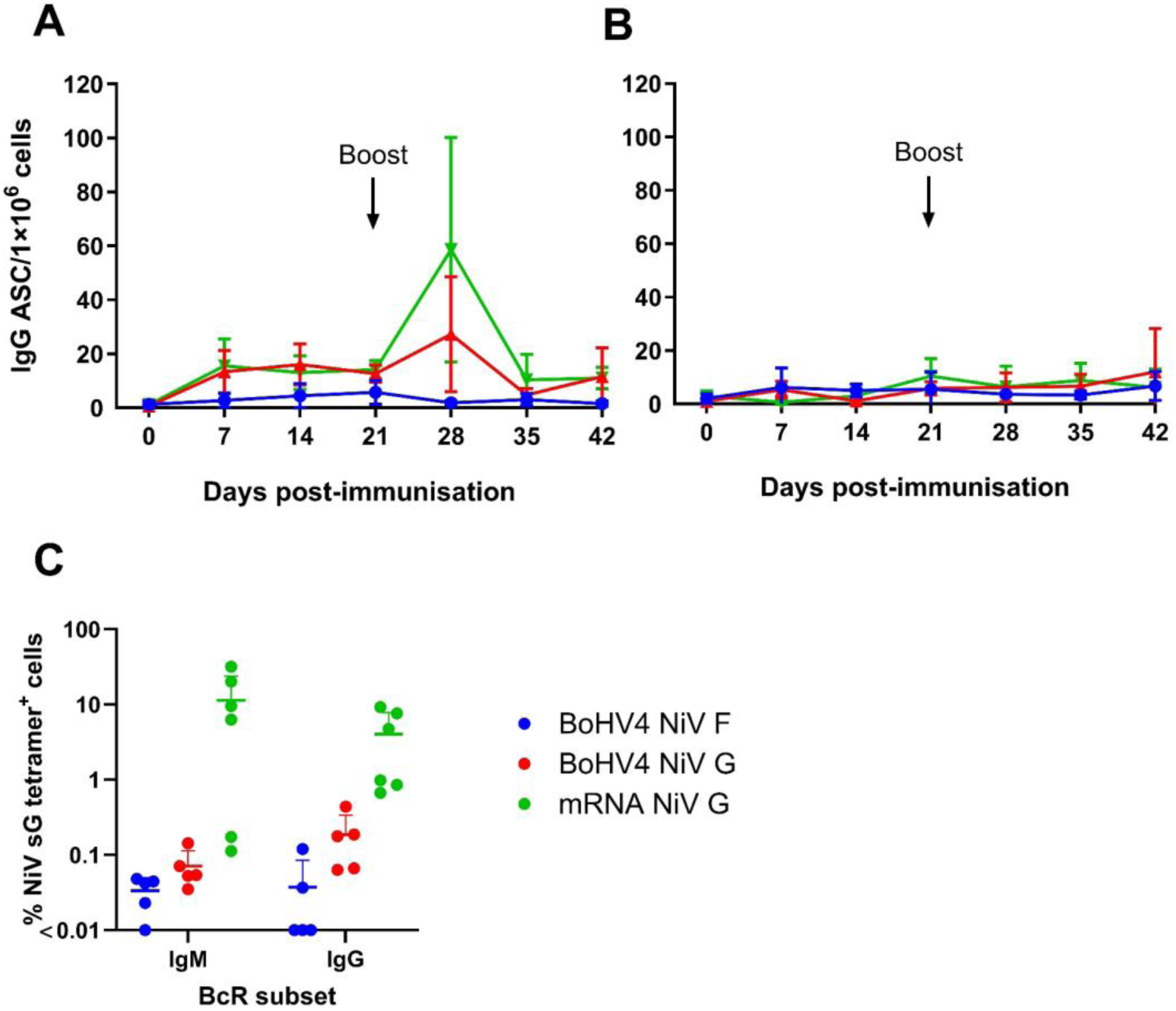
Evaluation of NiV-M G specific B cell responses in vaccinated pigs. Groups of pigs were immunised with BoHV-4 NiV F (n=5), BoHV-4 NiV G (n=6), or mRNA NiV G (n=6) on day 0 and boosted at day 21. IgG ELISpot assays were performed with PBMCs to enumerate plasma B cells secreting (A) NiV G or (B) KLH (irrelevant antigen) specific antibodies. Results are expressed as the mean number of IgG-antibody secreting cells (ASC) per million PBMC for each group and error bars represent the standard deviation (SD). (C) NiV-M G-specific IgM^+^ and IgG^+^ B cells on day 28 were assessed by tetramer labelling of previously cryopreserved PBMC and flow cytometry. Datapoints represent the % tetramer^+^ cells within the parent B cell population for each animal, horizontal bars represent the group median.

### Generation of recombinant porcine NiV G-specific mAbs

Aligned with the higher frequency of specific B cells within PBMC from the mRNA vaccinated group, three pigs were selected from this group on 28 dpv to perform single cell sorting of NiV-M sG-binding IgG^+^ B cells. From one of the plates (derived from a single donor pig), 25 out of 70 wells resulted positive for IgG1 heavy (IGHG1) chain PCR products, of which 13 were positive for the lambda light (IGL) chain and 17 for the kappa light (IGK) chain PCR products. Fifteen out of the 70 wells (21% of the total sorted cells) produced H and L chain pairs, while the remaining L or H chains remained unpaired. Of the 15 pairs (all gamma H chains IgG1), nine had an IGL chain and six had an IGK chain. The PCR products from the paired heavy and light chain gene segments were cloned into their respective expression vectors. Native pairs of H and L chain expressing plasmids were co-transfected to express 12 porcine recombinant mAbs. Of these, five (called A2, C1, D8, F4 and G9) bound to the NiV-M sG protein in ELISA (Supplementary Figure 4).

Amino acid sequences of complementarity determining regions (CDR) of H and L chains for the five porcine NiV G specific mAbs were analysed (Supplementary Figures 5 and 6). The CDRH3 lengths for mAbs C1, D8, F4 and G9 were average for pigs (12-14 amino acids), however, mAb A2 was amongst the longest ∼5% (20 amino acids) [45] (Supplementary Figure 6A). Analysis of level of somatic hypermutation (SHM) of gene segments for H and L chains was measured by distance in base pair from known germline sequences [46–48] (Supplementary Figure 6B). mAb F4 was the furthest from germline both in H and L chain V gene segments, whereas mAbs A2 and G9 were the closest to germline. The L chain sequences for mAbs A2 and G9 likely derive from the same *IGLV8-10* gene segment however, as the associated H chains are completely different, these mAbs are not derived from the same pre-immune clone. The IGK sequences for mAbs C1, D8 and F4 are distinct, although D8 and F4 are possibly derived from the same IGKV1 gene segment, whereas the C1 clone belongs to a different subgroup (the closest gene segment being *IGKV2-13*). None of the five IGHG1 clones completely matched known IGHV gene segments, which is not unexpected given the influence of SHM and allelic diversity. All the pig IGHV gene segments belong to the same subtype, and although CDRH1 and CDRH2 showed slight variations, they could not be reliably mapped to individual germline IGHV gene segments (Supplementary Figure 5).

### Assessment of binding and neutralising activity of porcine NiV G-specific mAbs

To more fully assess binding, titrations of the five porcine NiV G-specific mAbs were further evaluated for binding to recombinant NiV-M sG and HeV sG proteins by indirect ELISA (Figure 2A, B, respectively). All five mAbs specifically bound to NiV-M sG, however, only mAb A2 bound to HeV sG. Biolayer interferometry (BLI) was performed to determine kinetic constants for the interactions between mAbs and NiV-M sG (Figure 2C). All mAbs bound to NiV-M sG with high affinity; with Kd values < 0.1 nM.

**Figure 2.**
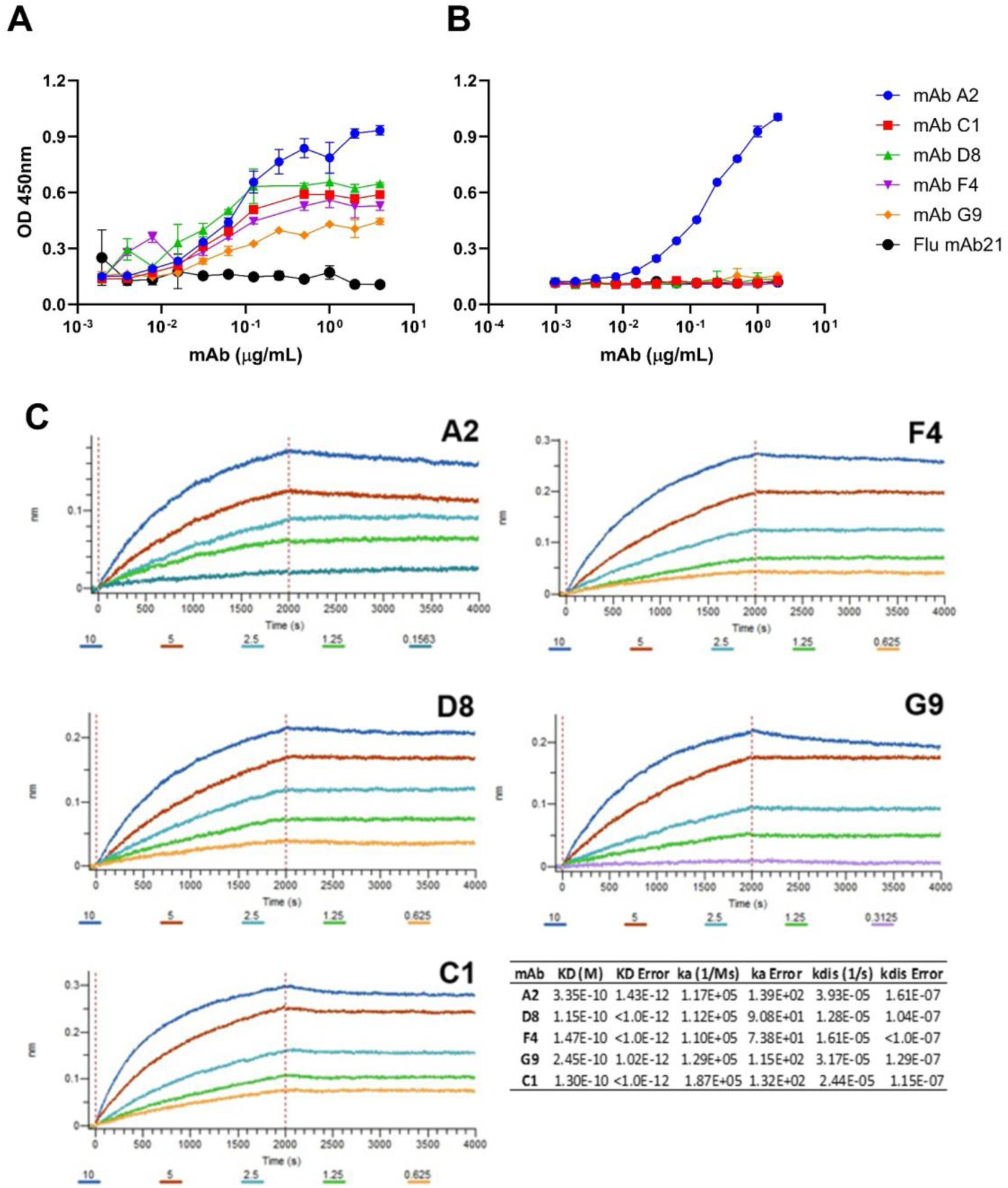
Assessment of the binding properties of NiV G specific porcine mAbs. Recombinant porcine mAbs were evaluated for binding to recombinant (A) NiV-M sG or (B) HeV sG by ELISA. An influenza A virus-specific porcine recombinant mAb (PB21) was used as negative control. Mean technical duplicate data ± SD from a representative experiment are shown. (C) BLI was conducted to determine mAb binding kinetics to NiV-M sG. Association and dissociation curves were determined for 2000 sec, to calculate Kd values, association and dissociation constants. Representative data from a single experiment are presented.

mAbs were next evaluated for their ability to neutralise NiV-Mpp and NiV-Bpp (Figure 3). Neutralising activity was compared to that provided by recombinant human mAb m102.4. All five porcine mAbs and mAb m102.4 neutralised NiV-Mpp (Figure 3A), whereas only mAbs A2 and m102.4 were able to neutralise NiV-Bpp (Figure 3B). Comparison of the calculated IC_50_ for each mAb, revealed no significant difference between mAb A2 and m102.4 in terms of their capacity to neutralise NiV-Mpp nor NiV-Bpp (Figure 3C). mAb C1 was the most potent neutraliser of NiV-Mpp with a significantly lower IC_50_ than all other mAbs (*p*<0.05), whereas mAb G9 was the weakest neutraliser presenting with an IC_50_ significantly higher than the other mAbs (*p*<0.05). Assessment of mAb neutralisation of NiV glycoprotein-mediated cell-cell fusion, showed three mAbs, A2, C1 and F4, capable of neutralising NiV-M fusion (Figure 3D), whereas only mAb A2 neutralised NiV-B fusion (Figure 3E). Comparison of the NiV-M fusion inhibiting titres (IC_50_) showed that mAb C1 was significantly more potent than the other two mAbs (*p* <0.01) (Figure 3F).

**Figure 3.**
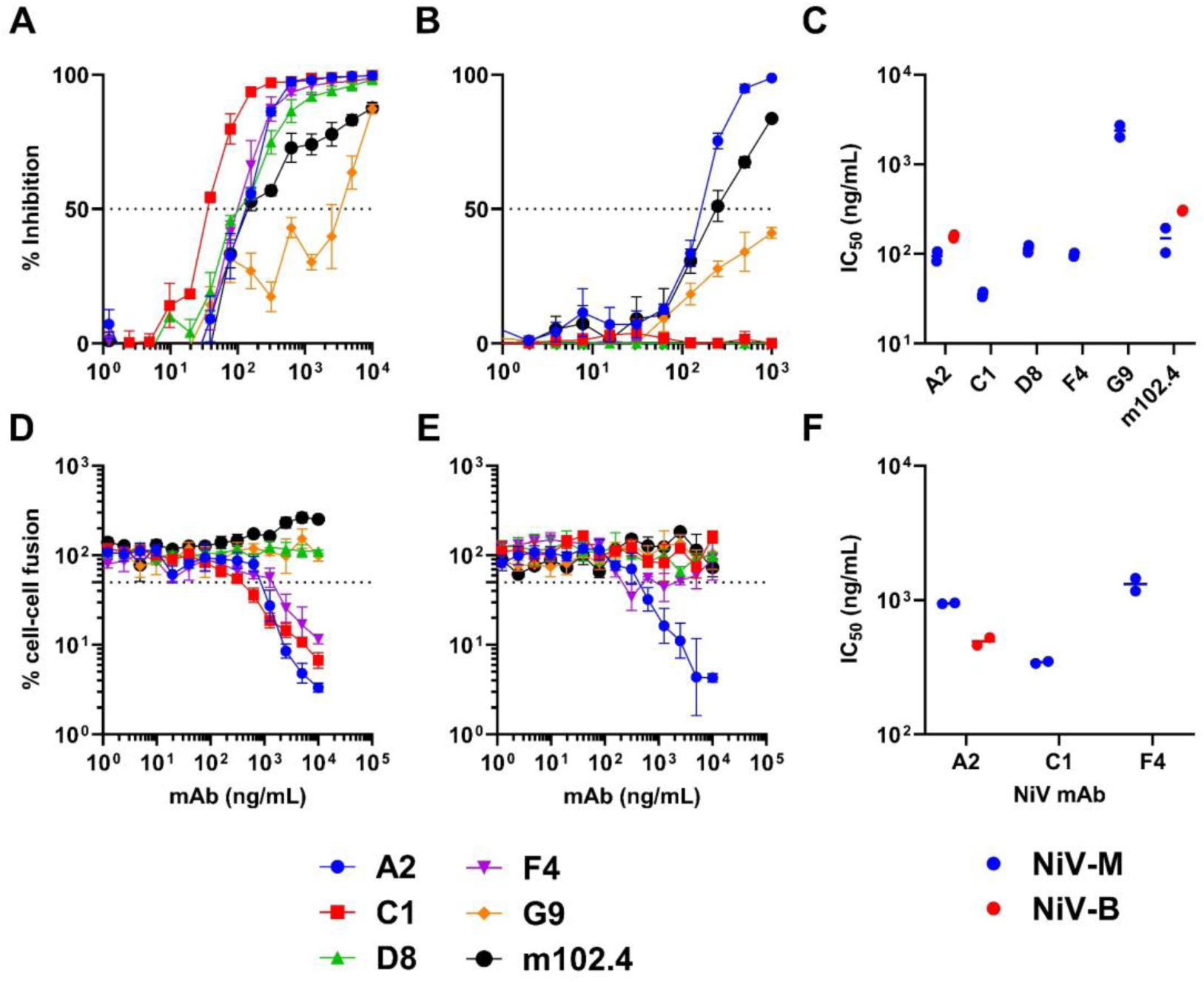
Assessment of the neutralising properties of NiV G specific porcine mAbs. Recombinant porcine mAbs were evaluated for neutralisation of (A) NiV-M or (B) NiV-B pseudovirus particles. Pseudovirus neutralisation titres were calculated as the inverse of the mAb concentration which showed a 50% inhibition of infection (IC_50_) (C). mAbs were also assessed for inhibition of (D) NiV-M and (E) NiV-B glycoprotein mediated cell–cell fusion; (F) with titres calculated as the inverse of the mAb concentration which showed a 50% inhibition of fusion (IC_50_) For A, B, D, and E, mean technical duplicate data ± SD from a representative experiment are shown. For C and F, each data point represents individual pig sera with lines denoting the median value.

### Elucidating epitopes targeted by NiV neutralising mAbs

#### Epitope binning by competition ELISA

To investigate whether the NiV G-specific porcine mAbs recognised overlapping epitopes, a competition ELISA was performed, which included mAb m102.4 as a reference. All mAbs were evaluated against each other in a reciprocal competition format using biotinylated mAbs. The percentage of residual binding was calculated after background correction, with values ≥90% interpreted as no competition. The resulting heatmap revealed distinct binding profiles among the mAbs (Figure 4). mAbs A2 and C1 showed minimal competition with mAb m102.4, indicating that these mAbs recognise non-overlapping epitopes and can simultaneously bind NiV sG. Additionally, mAb A2 showed no competition with mAb C1, suggesting recognition of a separate antigenic region. In contrast, mAbs D8, F4 and G9 showed reduced residual binding when assessed against m102.4, indicating partial or complete epitope overlap. These findings suggest the presence of multiple epitope groups on NiV-M sG and highlight mAbs A2 and C1 as interesting candidates for further characterisation of their binding sites, neutralising properties, and potential use in antibody combination strategies.

**Figure 4.**
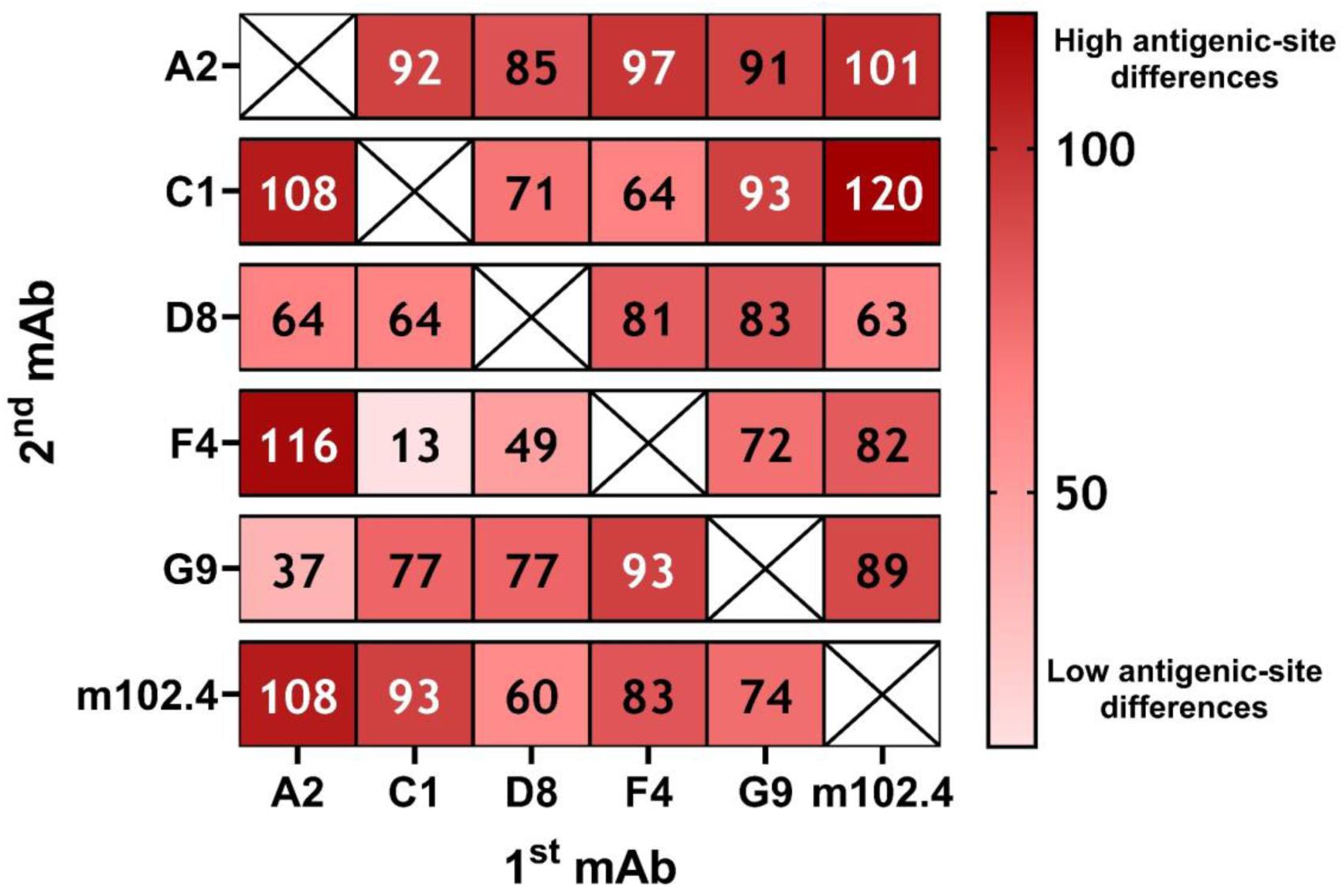
Competition binding profile of NiV sG-specific mAbs. Heatmap showing the reciprocal competition patterns among the mAbs determined by competition ELISA. The first-added unlabelled mAb (x-axis) was used as the competitor, while the second-added biotinylated mAb (y-axis) was used as the detection antibody. Values represent the percentage of residual binding after background correction, with higher values indicating minimal competition (non-overlapping epitopes) and lower values indicating increased epitope overlap. Crossed boxes indicate self-competition controls.

#### Structural determination of NiV G in complex with mAbs A2 and C1

To elucidate the molecular basis for targeting of NiV G by mAbs A2 and C1, recombinant Fabs were expressed and binding confirmed by ELISA (Supplementary Figure 7). We subjected a recombinantly-derived complex of NiV G RBD-Fab A2/C1 to single-particle cryo-EM analysis to yield a 3.19 Å resolution structure (Figure 5). The NiV G RBD forms a monomeric six-bladed β-propeller characteristic to receptor binding proteins (RBPs) of the *Paramyxoviridae* family [49]. Fabs A2 and C1 bind to the propeller at non-overlapping epitopes, providing a structural rationale for the non-competing nature of these two Fabs (Figure 4). For Fab A2, each loop of the CDR light chain and the CDRH3 of the heavy chain recognises blades β1 and β6 (defined as interface regions I1 and I2 in Figure 5G) and occludes ∼1,330 Å^2^ of the NiV G RBD surface and is stabilised by four hydrogen bonds (Figure 5B, 5C, 5E, 5G). Interestingly, this epitope includes residues involved in henipaviral RBP dimerisation, as identified in previous structural investigations [50,51] (Supplementary Figure 8). Fab C1 recognises blades β2 and β3 of the NiV G RBD β-propeller (defined as interface regions I3 and I4 in Figure 5G) in an interaction stabilised by fourteen hydrogen bonds and one salt bridge (Figure 5B, 5D, 5F, 5G). All CDR loops from Fab C1 are engaged in the interaction and occlude approximately 1,700 Å^2^ of the RBD surface (Figure 5D, 5F). Interestingly, neither of the Fab A2 or Fab C1 epitopes overlap with the receptor-binding site (Figure 5), indicative that the Fabs do not directly impede ephrinB2/B3 recognition. Thus, the mAbs likely act either by sterically impeding receptor recognition through their higher-order structure of a full-length antibody or by interfering with fusion activation downstream of host cell receptor engagement.

**Figure 5.**
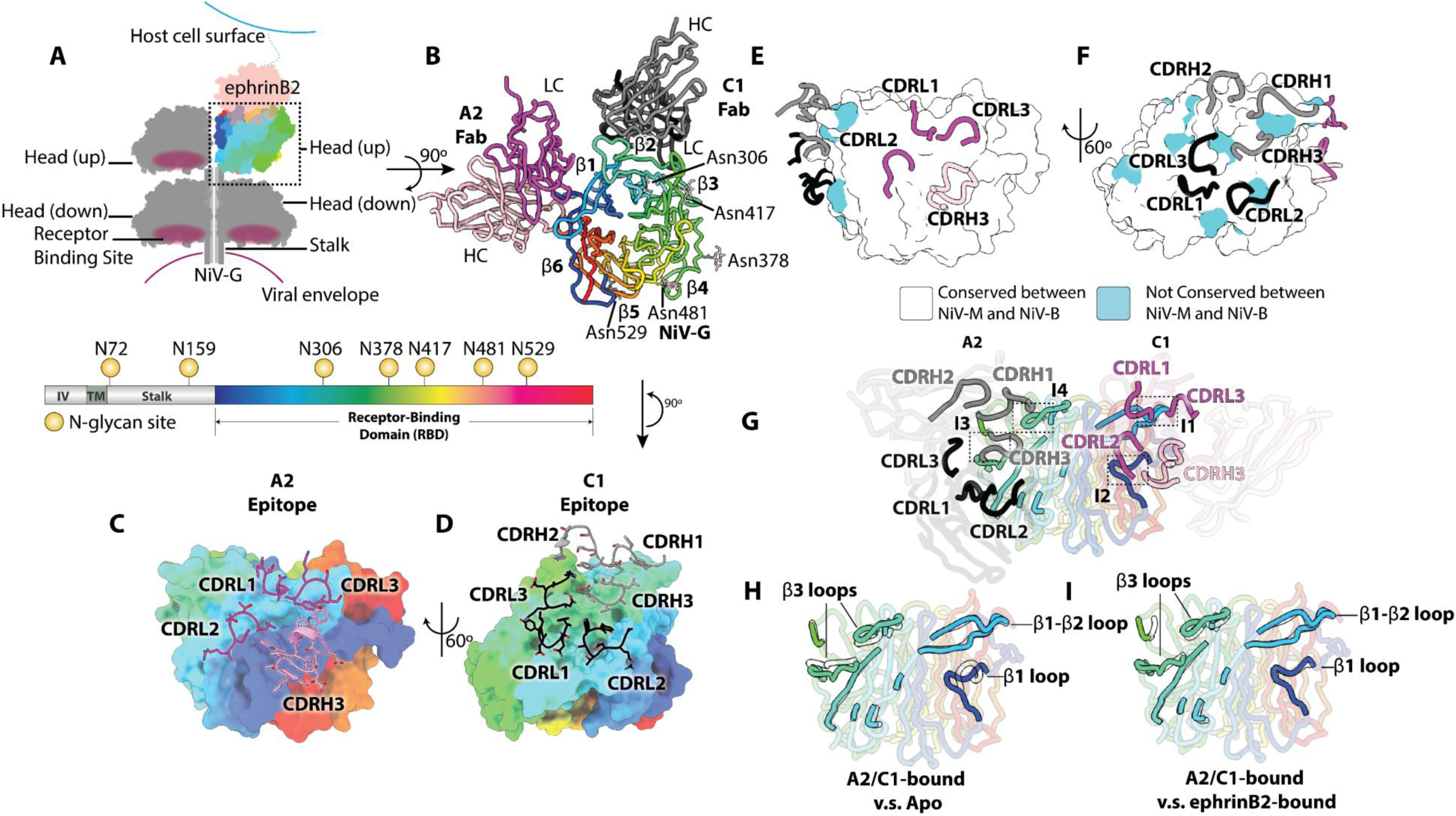
Structural characterisation of NiV G RBD in complex with porcine Fabs A2 and C1. (A) Diagram displaying native interaction between the NiV G tetramer [51] and ephrinB2 on the host cell surface. Only one RBD (head domain) is engaged with ephrinB2 in this diagram, which is coloured with rainbow from the N- (blue) to C- (red) terminus. Below is a domain plot showing NiV G domain boundaries and predicted N-linked glycosylation sequons (NXT/S, where X does not equal P). IV = intra-viral, TM = transmembrane. The domain plot was created with the IBS 2.0 server [69]. (B) Top-down view looking at the NiV G-FabA2/FabC1 complex structure. β1-β6 annotations indicate the positions of β-propeller blades. (C) Contacts between Fab A2 CDR loops with the surface of NiV G. CDR loops of the light chain are coloured magenta and CDR loops of the heavy chain are pink. Amino acid sidechains of CDR loops are shown as sticks. (D) C1 epitope mapped onto the NiV G RBD surface. CDR loops of light chain are coloured with black and those of heavy chain are coloured with grey. Sticks representation of the amino acid sidechains is shown for CDR loops. (E) and (F) Sequence conservation between NiV-M and NiV-B on the RBD surface. CDR loops are presented as tubes with the same colouring as (C) and (D). (G) A view looking at the junction between β1 and β2 showing both A2 and C1 epitopes with the same colouring as (B)-(F). Interface regions (I1-I4) are highlighted. (H) An overlay of A2 and C1 epitopes in Fabs A2/C1-bound structure and the apo structure (PDB: 2VWD) [34]. Non-interacting residues are transparent. The NiV G apo structure is presented with transparent tubes with sharp black outline. (I) An overlay of A2 and C1 epitopes in Fabs A2/C1-bound structure and the ephrinB2-bound structure (PDB: 2VSM) [33]. Non-interacting residues are transparent. The ephrinB2-bound NiV G structure is presented with transparent tubes with sharp black outline.

### Molecular specificity of mAb A2 and C1 for NiV G

Overall, NiV G RBD structures of apo and ephrinB2-bound states both exhibit an overall ∼0.9 Å root-mean squared deviation (rmsd) from the A2/C1-bound state (over 408 and 411 aligned Cα atom pairs, respectively). Interestingly, when focusing on the epitopes, structural overlay analysis reveals that A2/C1-bound NiV G most closely resembled the ephrinB2-bound state [33], when compared to the apo NiV G state [34] (Figure 5G-I). Indeed, residues comprising the A2 epitope region exhibit 2.1 Å rmsd from the apo state (21 Cα atom pairs) compared to only 0.4 Å rmsd for the ephrinB2-bound structure (20 Cα atom pairs). Similarly, the conformation of residues comprising the C1 epitope are structurally more similar to the ephrinB2-bound state (0.8 Å rmsd over 31 Cα atom pairs) than apo state (1.3 Å over 31 Cα atom pairs).

The interaction between region I1 and Fab A2 relies mostly on hydrophobic interactions and π-stacking between NiV G residue in the β1 blade, where Phe266, Arg51, and Trp103 on the β1-β2 loops interact with residues in the Fab CDRH3 loop, respectively (Supplementary Figure 9). Additionally, CDRH3 loop residue, Asp111, forms a hydrogen bond with NiV G Ser204 inducing this residue to orient away from the RBD in a conformation similar to that observed in the ephrinB2-bound state [33] (Supplementary Figure 9-10).

Fab C1 recognises an epitope near the β3 blade (defined as interface regions I3 and I4 in Figure 5G). Indeed, recognition breaks the NiV G Arg338−Glu422 salt bridge, which naturally forms in the apo form and induces Arg338 to point outwards from the RBD centre, similar to that observed in ephrinB2-bound state [33] (Supplementary Figure 9). This interaction includes multiple hydrogen bonds between Tyr33/Ser52/Tyr59 on the CDRH loops and Glu422 on the RBD, Ser98 on the CDRL3 loop and Asn423 on the RBD, and CDRL1 loop mainchain and Arg338, and a π-stacking interaction between Phe36 on the CDRL1 loop and Arg338 (Supplementary Figure 9). This complex interaction is further stabilised by a hydrophobic patch comprising of CDRH1 and CDRH3 at interface region I4.

Our data also provide a structural rationale for why mAb A2 cross-reacts with the NiV-B G, whereas mAb C1 does not. While the 21 residues comprising the A2 epitope are fully conserved between NiV-M and -B, we observed 5 (*i.e.*, 5 out of 31 residues) sequence differences at the C1 epitope: S325N, G328E, G329S, L335F, and S339N (Supplementary Figure 11). The most pronounced of these differences are G328E and G329S, where the introduction of glutamic acid and serine residues in this region of the interface are less compatible with the hydrophobic nature of this region of the interface (Supplementary Figure 9).

### Evaluation of the protective efficacy of mAb A2 in vivo

To determine the effectiveness of mAb A2 as a candidate therapeutic, a comparative challenge study was performed with mAb m102.4. Extensive preclinical studies have been performed utilising mAb m102.4 demonstrating protection against NiV-M challenge in ferrets [4] and HeV challenge in African green monkeys [52]. In addition, m102.4 has been used in compassionate use situations to individuals following a high exposure risk [53]. Here, hamsters were administered with mAb A2, mAb m102.4 or a combination of the two mAbs, 1 day prior to challenge with NiV-B (Figure 6). All animals inoculated with mAb m102.4 were fully protected following challenge. Animals that received mAb A2 exhibited a 60% survival rate, while a combination of both mAbs m102.4 and mA2 resulted in complete protection. All challenge control animals lost weight post-challenge and met humane endpoints requiring euthanasia between 3-5 days post infection. NiV-B shedding was measured at 3 and 5 dpi (Figure 6C). Infectious NiV-B titres were detected in 4/6 of the challenge control animals, compared to a single animal in the mAb m102.4 and mAb combination groups, and 2 animals in the mAb A2 treated groups. Only the m102.4 treatment group showed a statistically significant reduction in oral swab titres (*p* < 0.05). At 5 dpi, two of the three surviving challenge control animals showed infectious NiV-B in oral swabs and no titres were detectable in the three mAb treatment groups. NiV-B titres were only measurable in nasal swabs from one challenge control animal and two mAb A2 treated animals at 3 dpi. NiV-B was undetectable in rectal swab samples in all groups at both timepoints.

**Figure 6.**
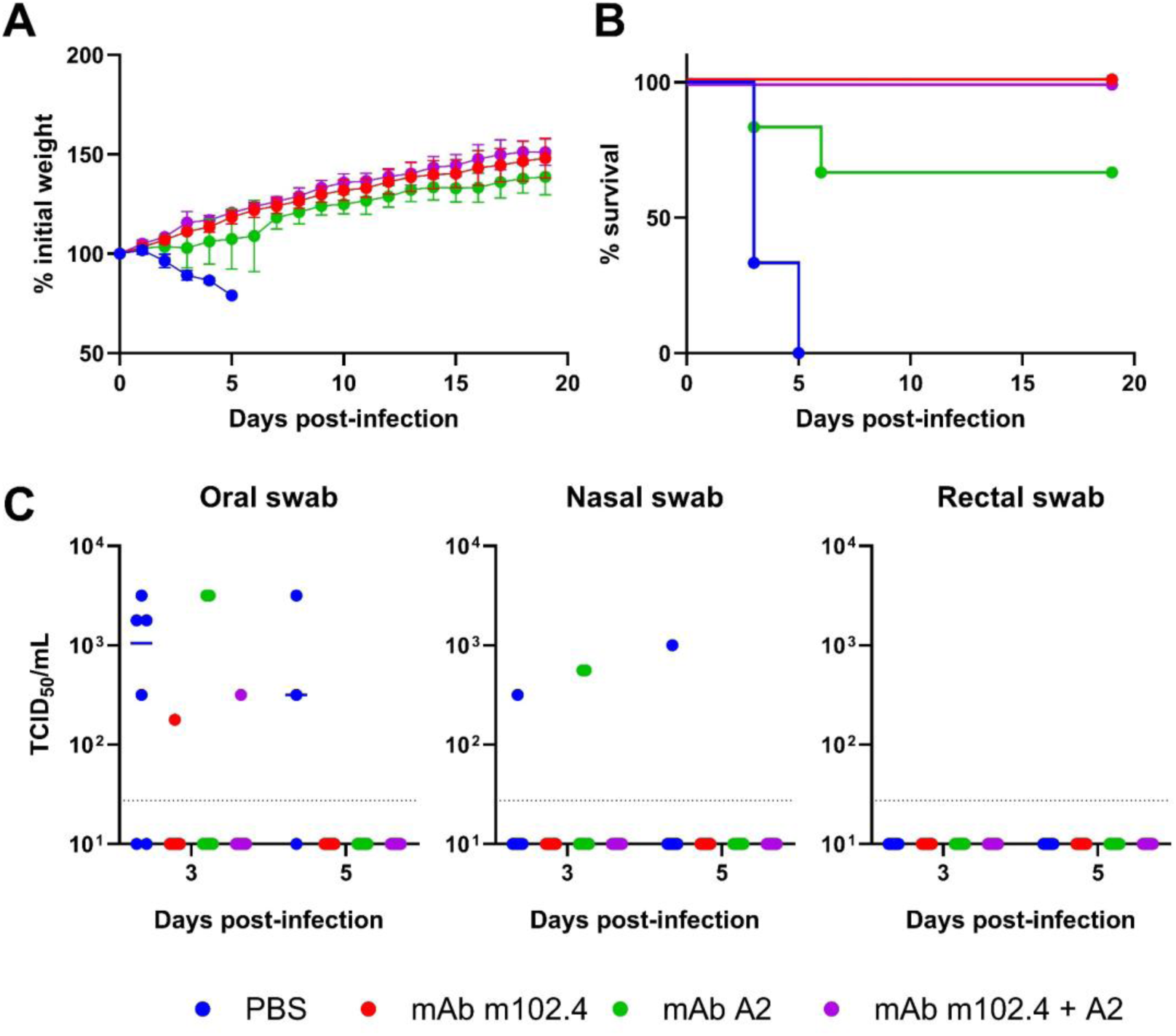
Assessment of the protective efficacy of mAb A2 alone and in combination with mAb m102.4. Groups of hamsters (n=6) were inoculated with PBS, mAb m102.4, A2, or a mixture of m102.4 and A2 (0.5 mg/mAb), one day prior to NiV-B challenge. Animals were (A) weighed (group mean ± SD are shown) and (B) clinical signs scored daily to assesses severity of disease and euthanised when humane endpoints were met. (C) Oral, nasal, and rectal swab samples were collected 3 and 5 dpi and infectious NiV-B titres determined by TCID_50_ assay. Datapoints represent individual animals and bars show the group median value.

## Discussion

NiV poses a significant epidemic threat due to the widespread distribution of fruit bat reservoir species across South and Southeast Asia, coupled with the density of pigs and other livestock species that can act as intermediate amplifying hosts. NiV infection in humans results in severe respiratory and neurological disease with a case fatality rate estimated to be ∼80% [54]. Currently, there are no specific treatments for NiV infection, which is instead limited to supportive medical care. NiV neutralising mAbs are considered a promising route to effective post-exposure or early treatment therapies. mAbs targeting either the F or G glycoproteins can effectively neutralise NiV and have been shown to protect against severe NiV or HeV infections in preclinical animal models [4–6,9–11]. The leading candidate, m102.4, targets the NiV/HeV G RBD, preventing engagement with the ephrinB2/B3 receptors on host cells [55,56]. m102.4 has been used to treat NiV infected patients on a compassionate basis and has successfully passed a phase 1 clinical trial [7]. Whilst the NiV G RBD must be structurally constrained, to an extent, to remain functional, *in vitro* passage of NiV and HeV in the presence of m102.4 led to the generation of escape mutant viruses after only three passages [55]. Consequently, recent efforts have been focussed on novel mAb combinations targeting discrete epitopes, which may act synergistically and reduce the risk of virus escape from treatment [12,27]. This study aimed to contribute to these efforts through the characterisation of NiV-neutralising mAbs from immunised pigs.

Pigs can play a key role in NiV epidemiology as exemplified by their role in the first, and still most severe, outbreak in Malaysia and Singapore [3]. In addition to being a target species for a NiV vaccine, pigs also provide a large, outbred animal model for evaluating NiV vaccine candidates being developed for humans [57,58]. mAbs can also be isolated from immunised/infected pigs, using single cell technologies, which resemble antibodies observed in humans [22,23]. Taking advantage of an immunogenicity study in pigs evaluating the immunogenicity of NiV G or F delivered using mRNA or BoHV-4 vectors [16,20], we sought to isolate mAbs from NiV G binding B cells. Due to the more potent plasma cell response post-boost, B cells were sorted from mRNA pigs. A panel of five recombinant mAbs were produced, which all bound NiV-M sG with sub-nanomolar affinity. That only mAb A2 cross-reacted with HeV sG was unsurprising, since the serum from the donor animal presented with a high NiV-M neutralising titre (ND_50_ 2560) but low HeV neutralising titre (ND_50_ 10) [20]. However, since the NiV-B neutralising titre (ND_50_ 640) in serum was less than a log_10_ lower than NiV-M, it was more surprising that only mAb A2 was able to neutralise NiV-B. The four strain specific mAbs were unique, differed in their neutralising potency and targeted more than a single antigenic site. Interpretation of this apparent predominance of strain-specific mAbs is limited by the small number isolated from a single animal. Nevertheless, our data contrast with 25/27 mAbs, isolated from immunised mice using NiV-M sG as bait [59], neutralised both NiV-M and NiV-B [60]. mAbs A2 and C1 were selected for structural analysis since they showed minimal competition with each other and m102.4, suggesting they recognise discrete epitopes and simultaneously bind NiV G. Single particle cryo-EM of NiV-M G RBD complexed to Fabs A2 and C1 confirmed distinct epitopes that did not overlap with the receptor-binding site, targeted by m102.4. This suggests that neutralisation may occur through mAb binding resulting in steric impedance of receptor binding or via interference with fusion activation following receptor engagement. Interestingly, the A2/C1-bound NiV G RBD most closely resembled the ephrinB2-bound state [33], which could suggest conformational trapping of G in a receptor-bound-like, fusion-incompetent state, preventing activation of F protein, and thereby blocking membrane fusion. This hypothesis is supported by the observation that both mAbs blocked NiV glycoprotein mediated fusion in a fusion inhibition test [31]. Epitopes targeted by NiV G-specific mAbs have previously been mapped and binned [4–6,51,55,59–64] (Figure 7 and Supplementary Table 2). Out of 46 mAbs identified in the literature, 21 mAbs, including m102.4, bind to the receptor-binding site, 17 mAbs bind near the stalk on the RBD, five mAbs, including A2, bind to the dimerization interface, one bind near blade β1 close to receptor-binding site (S2B10), one bind to blade β3 (S1E2), and only C1 binds to the region connecting blades β2 and β3. The surface region near the β2 and β3 blades, however, exhibits a lower level of sequence conservation between NiV-M and NiV-B. Thus, mAbs such as C1, which are directed against this region, are less likely to cross-neutralise NiV-B. Blades β4 and β5 are largely untargeted. This may be due to the higher density of N-linked glycans, which may mask this region during antigen presentation [65] (Figure 7). The epitopes bound by mAbs A2 and C1 therefore represent less commonly targeted regions on the NiV G surface. mAbs targeting these regions may likely function synergistically with antibodies that target the more immunodominant ephrin-B2 binding site.

**Figure 7.**
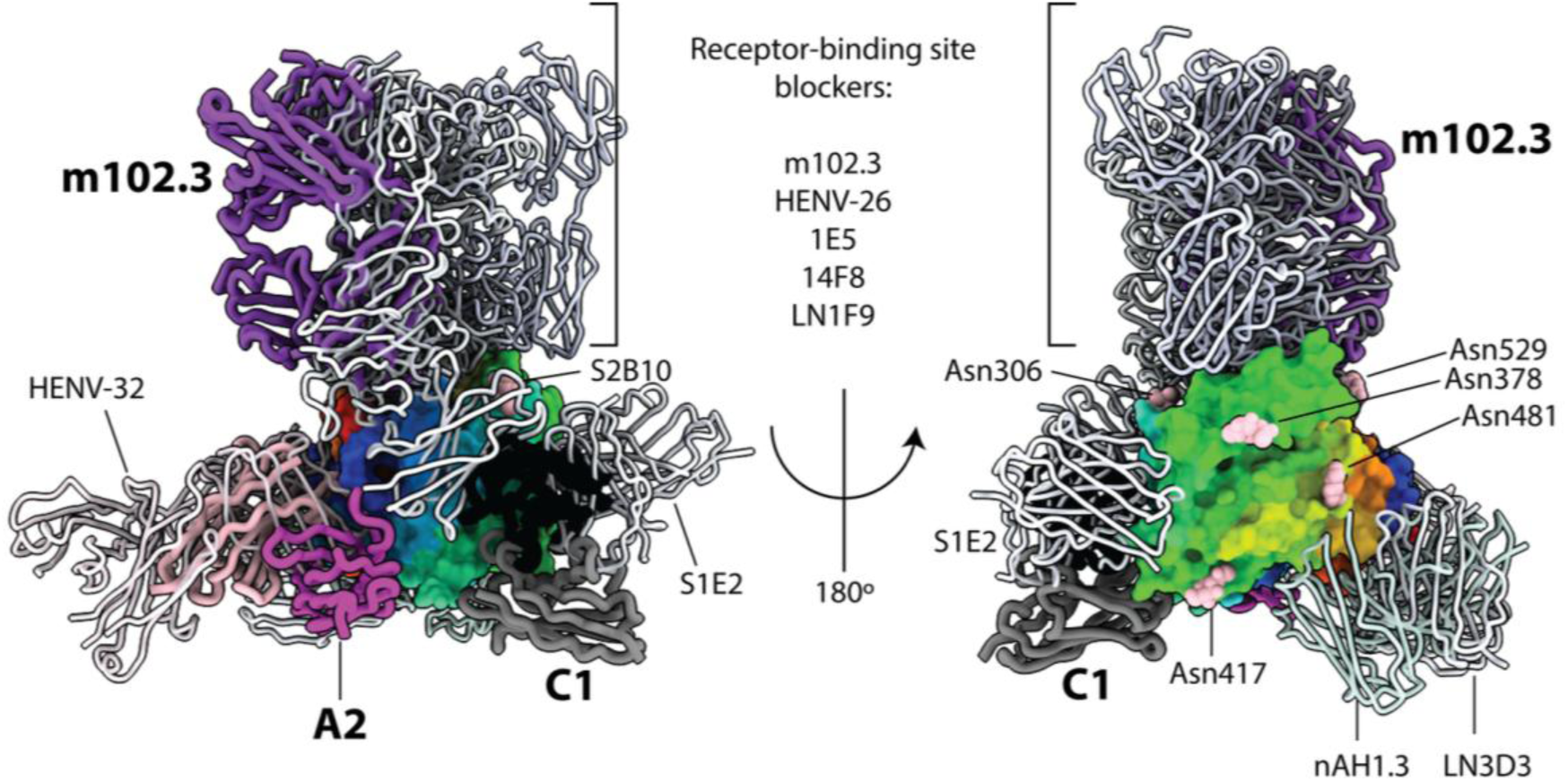
Antigenic landscape of the NiV G RBD. Representative mAbs structures from the literature (Supplementary Table 2) are presented as thin cartoon tubes and coloured grey except mAb m102.3, which targets the same epitope as m102.4, is represented as a purple thick cartoon tube. mAbs A2 and C1 are represented by thin cartoon tubes coloured as in Figure 5B. NiV G RBD is represented by a surface rainbow coloured from N-terminus (blue) to C-terminus (red), decorated with N-acetylglucosamine residues as pink spheres.

mAb A2 and m102.4 showed comparable neutralising potency *in vitro*. However, A2 was less effective than m102.4 in protecting hamsters against disease following NiV-B challenge. The reasons for this remain unclear. In addition to virus neutralisation, mAb-mediated protection may be reliant on Fc-mediated effector functions [66], which include activation of complement, phagocytes and NK cells. Human IgG subclasses are reported to bind to murine FcγRs with broadly similar affinity to orthologous human receptors [67,68]. However, the interactions between human and porcine IgG and cricetine FcγRs remain undefined and this may, in part, explain the difference in efficacy between m102.4 and mAb A2. Future studies could compare the efficacy of humanised mAb A2 and further explore the benefits of mAb combinations at reduced dosages.

In conclusion, vaccination of pigs with mRNA vectored NiV sG elicited robust antigen-specific B cell responses that enabled the isolation of high-affinity mAbs. Functional and structural analyses identified two potent neutralising mAbs, A2 and C1, which recognised distinct epitopes outside the canonical ephrinB2/B3 receptor-binding site. Despite not directly competing with receptor engagement, both mAbs neutralised NiV and stabilised conformations of NiV G that more closely resembled the receptor-bound state than the apo structure, suggesting disruption of the conformational transitions required for fusion activation. mAb A2 cross-neutralised both NiV-M and NiV-B, cross-reacted with HeV sG, and provided partial protection against lethal NiV-B challenge *in vivo*, while combined administration of A2 and m102.4 provided complete protection. These findings contribute to our understanding of the NiV G antigenic landscape, and the development of mAb combinations, that exert complementary mechanisms of neutralisation for therapeutic intervention against henipavirus infection.

## Author contribution

CRediT: **Miriam Pedrera**: Investigation, Methodology, Visualisation, Writing – original draft; **Nichakorn Pipatpadungsin**: Data curation, Formal analysis, Investigation, Methodology, Visualisation, Writing – original draft; **Darwyn Kobasa**: Investigation, Methodology, Visualisation, Writing – review & editing; **Ahmed M.E. Elrefaey**: Investigation, Methodology, Visualisation, Writing – original draft; **Barbara Holzer**: Methodology, Visualisation, Writing – review & editing; **Rebecca K. McLean**: Funding acquisition, Investigation, Writing – review & editing; **Bryce Warner**: Investigation, Writing – review & editing; **Robert Vendramelli**: Investigation, Writing – review & editing; **Nazia Thakur**: Investigation, Writing – review & editing; **Robert Stass**: Investigation, Writing – review & editing; **Jack W. P. Hayes**: Investigation, Writing – review & editing; **Lobna Medfai**: Investigation, Writing – review & editing; **Joshua E. Sealy**: Investigation, Methodology, Writing – review & editing; **Sylvia Crossley**: Investigation, Methodology, Writing – review & editing; **John C. Schwartz**: Formal analysis, Visualisation, Writing – review & editing; **Danish Munir**: Investigation, Methodology, Writing – review & editing; **William Mwangi**: Funding acquisition, Investigation, Methodology, Writing – review & editing; **Dalan Bailey**: Funding acquisition, Supervision, Writing – review & editing; **Thang Truong**: Investigation, Writing – review & editing; **Elma Tchilian**: Funding acquisition, Supervision, Writing – review & editing; **Bradley Pickering**: Funding acquisition, Investigation, Methodology, Supervision, Writing – original draft; **Thomas A. Bowden**: Funding acquisition, Supervision, Writing – review & editing; **Simon P. Graham**: Conceptualisation, Funding acquisition, Project administration, Supervision, Writing – original draft.

## Acknowledgements

We thank the Animal Sciences and Pathology Department, Animal and Plant Health Agency, UK for the care of animals and provision of samples; Gaetano Donofrio, University of Parma, Italy, for providing the BoHV-4 vectored vaccine candidates; Norbert Pardi and Drew Weissman, University of Pennsylvania, USA for providing the mRNA vectored vaccine candidate; Jamie R. M. Webster and the University of Birmingham Protein Expression Facility, UK, for the expression and purification of NiV sG protein; Rüdiger Raue and Mercedes Mourino, Zoetis for providing the HeV sG protein; Ariel Isaacs, Keith Chappell and Daniel Watterson, University of Queensland, Brisbane, Australia, for the m102.4 expression plasmids; Chris Broder, Uniformed Services University, USA, for allowing studies with m102.4 (which is now licensed to Mapp Biopharmaceutical); and The Pirbright Institute Flow Cytometry Science Technology Platform for assistance with cell sorting. We acknowledge Diamond Light Source for access and support of the cryo-EM facilities at the UK national Electron Bio-Imaging Centre (eBIC), proposal BI34631.

## Data availability statement

All data supporting the findings of this study are included within the article and its supplemental online material. New sequence data is publicly available: Recombinant NiV-M G-specific porcine monoclonal antibodies - GenBank accessions mAb A2 IGHG1 PZ831670 and IGL PZ831671, mAb C1 IGHG1 PZ831672 and IGK PZ831673, mAb D8 IGHG1 PZ831674 and IGK PZ831675, mAb F4 IGHG1 PZ831676 and IGK PZ831677, and mAb G9 IGHG1 PZ831678 and IGL PZ831679, and structural data: EMDB entries EMD-59475, and PDB 33QO. The original data are available from the corresponding author, SPG, upon reasonable request.

