## Supplemental Materials for "Isolation and characterisation of Nipah virus neutralising candidate therapeutic monoclonal antibodies from an mRNA-immunised pig"

### Supplementary Material

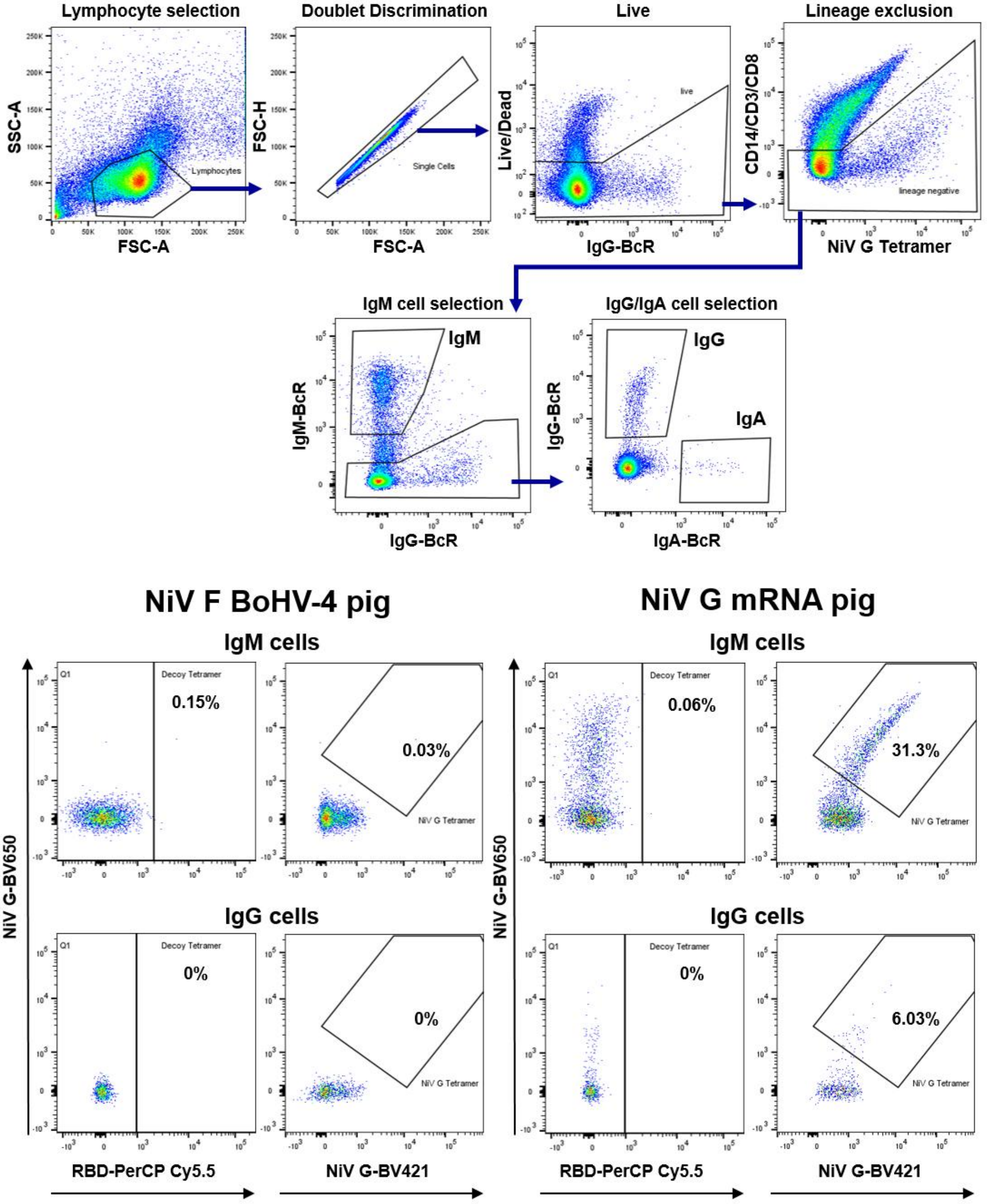


Supplementary Figure 1. Flow cytometric gating strategy to assess NiV G tetramer staining of porcine B cells. Representative data shown are from a BoHV-4 NiV F and an mRNA NiV G immunised pig at 28 days post-immunisation (7 days post-boost).

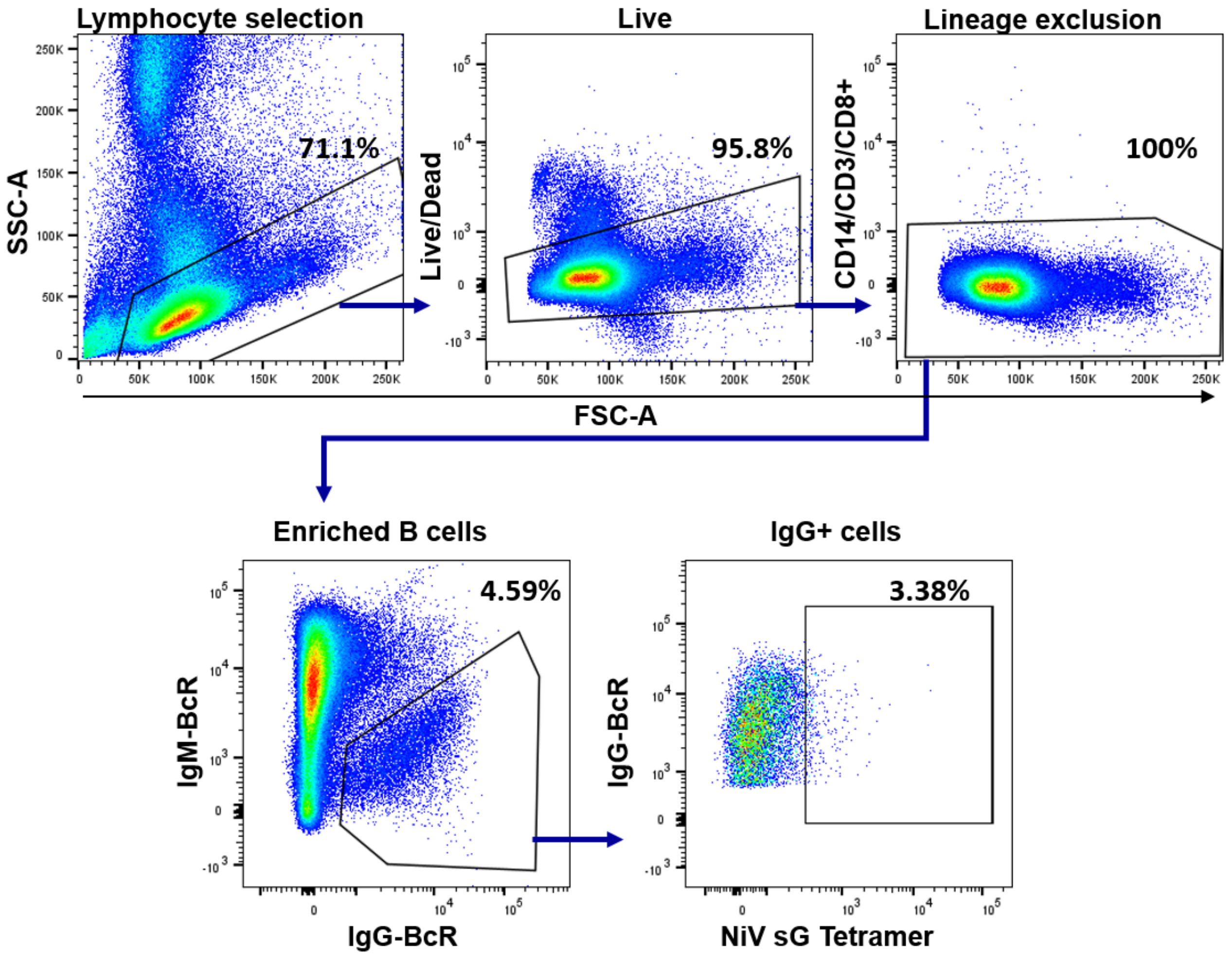


Supplementary Figure 2. Flow cytometric gating strategy to sort NiV sG tetramer binding enriched IgG$^{+}$ porcine B cells. Representative data shown are from an mRNA NiV G immunised pig 28 days post-immunisation.

Supplementary Table 1. Cryo-EM data collection and refinement statistics for NiV G RBD–Fab A2/Fab C1.

| | **NiV-G–FabA2/FabC1** |
|---|---|
| **Data collection and processing** | |
| Voltage (kV) | 300 |
| Electron exposure ($e^-$/ Å$^2$) | 50.00 |
| Defocus (μm) | 1.5 – 3.3 |
| Pixel size (Å/pix) | 0.831 |
| Symmetry | C1 |
| Final particle images (no.) | 119,065 |
| Map resolution (Å)[a] | 3.19 |
| FSC threshold | 0.143 |
| **Refinement** | |
| Initial model used | ModelAngelo[1] |
| Model resolution (masked) (Å) | 3.1/3.1/3.4 |
| FSC threshold | 0/0.143/0.5 |
| Map sharpening B factor (Å$^2$) | -135.5 |
| Model composition | |
| Non-hydrogen atoms | 6,834 |
| Protein residues | 876 |
| Ligands | 12 |
| B factors (Å$^2$)[b] | |
| Protein | 28.7/143.0/70.0 |
| Ligand | 69.0/161.7/111.3 |
| R.m.s deviations[c] | |
| Bond lengths (Å) | 0.004 |
| Bond angles (°) | 0.698 |
| **Validation** | |
| MolProbity score[d] | 2.63 |
| Clashscore | 9.73 |
| Rotamer Outliers (%) | 5.4 |
| Ramachandran[e] | |
| Favoured (%) | 89.4 |
| Allowed (%) | 10.6 |
| Outliers (%) | 0.00 |

[a]Gold standard FSC=0.143 from cryosparc v5.0.6 [2].

[b]B factors are listed as min/max/mean.

[c]RMS deviations: root mean square deviation from ideal geometry.

[d/e]Ramachandran analysis determined with the Molprobity server [3].

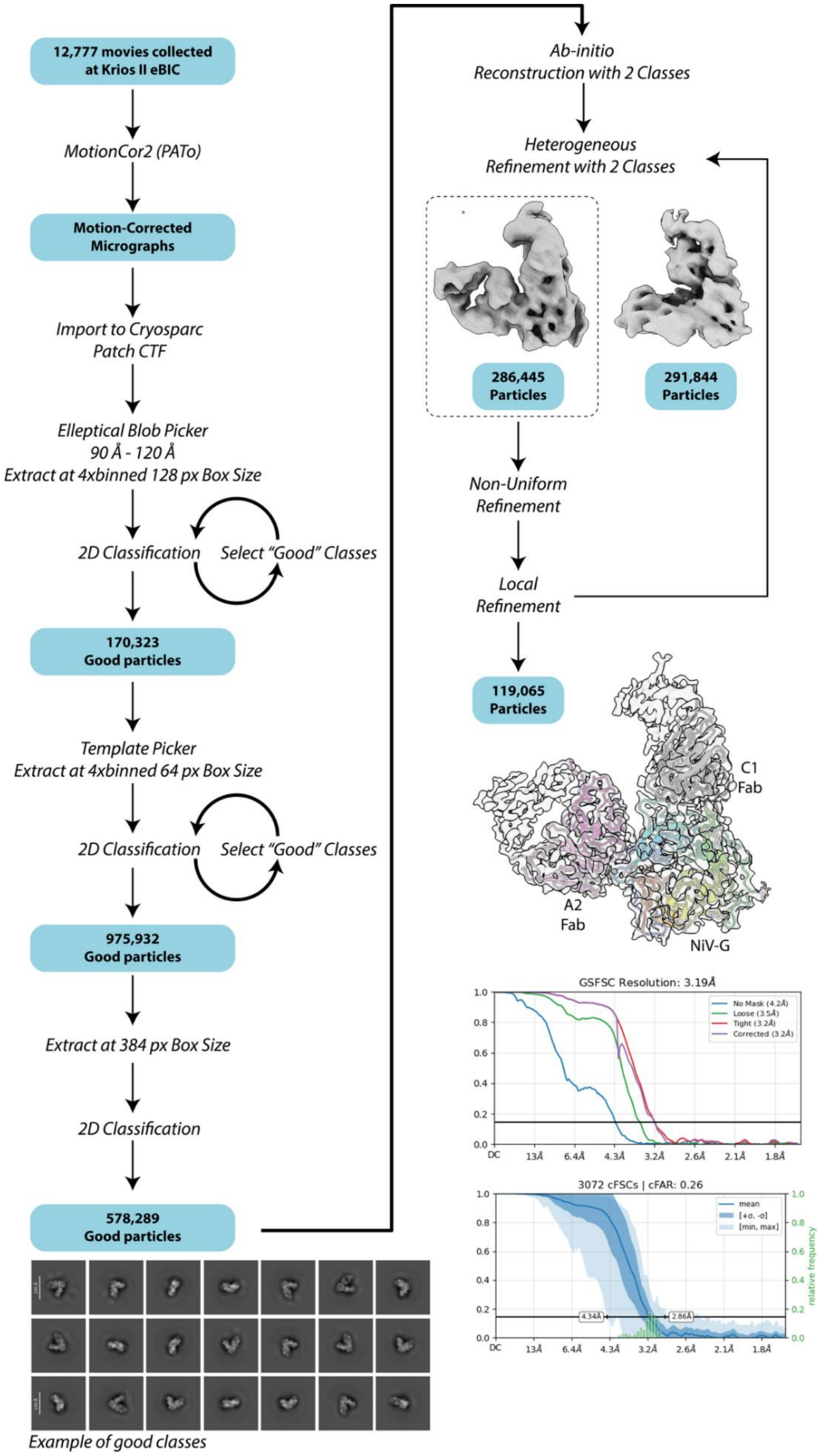


Supplementary Figure 3. Cryo-EM data processing workflow. Motion-correction was performed with MotionCor2 [15] in the PATo autoprocessing pipeline [16]. Main processing was performed in Cryosparc v5.0.6 [2]. Gold-standard Fourier-shell correlation (GSFSC) and conical Fourier-shell correlation (cFSC) plots with cFSC area ratio (cFAR) reported from the final local refinement [2] are also shown.

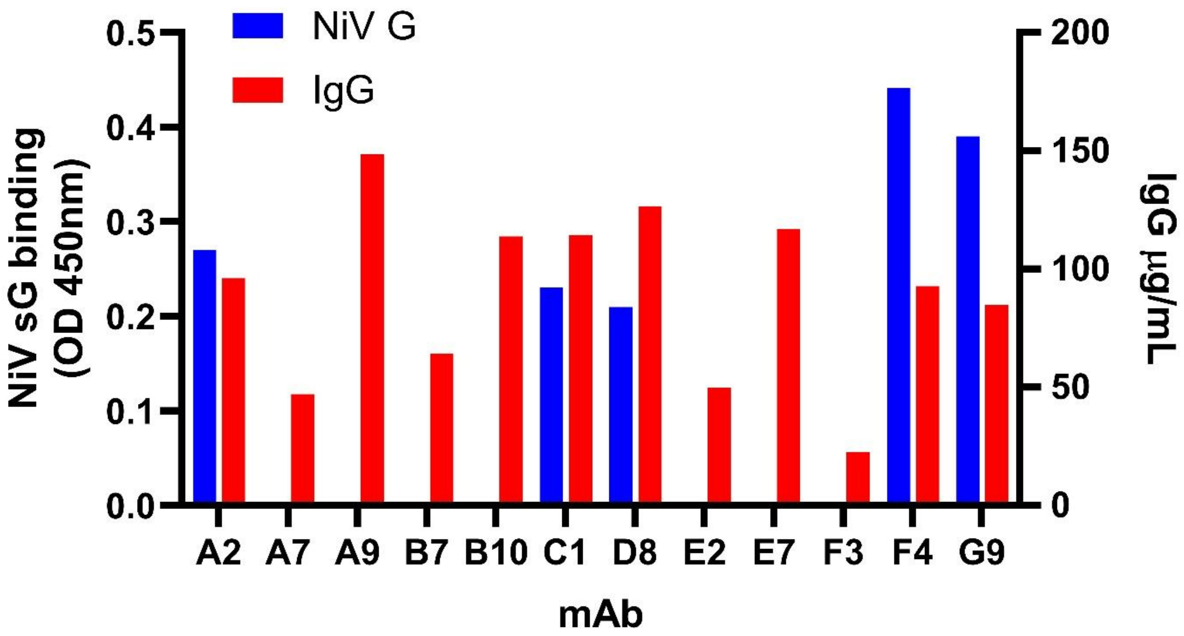


Supplementary Figure 4. Screening of the recombinant porcine mAbs by ELISA. Porcine IgG ELISA was used to measure mAb concentration (µg/ml) (right y-axis), and NiV sG ELISA was used as an initial assessment of antigen-specific binding (left y-axis).

Amino acid sequences. L. CDR1. CDR2. CDR3. C. conserved motifs/residues

>IGHG1_A2
…LSVLLMGCVAEEKLVESGGGLVQPGGSLRLSCVGSGFTFSSTYINWVRQAPGKGLEWLAAISTSGGSTYYADSVKGRFT
ISRDNSQNTAYLQENSLRTEDTARYYCARGIETWGETDWAPDYPMDLWGPGVEVVVSSAPKTAPSVYPLAPCGRDTSGPN
VALGCLASSYFPEPVTMTWNSGALTSGVHTFPSVLQPSGLYSLSSMVTVPASSLSS…

>IGL_A2
…LSVLLMGCVAQTVIQEPAMSVSPGGTVTLTCAFSSGSVTAPNYPSWFQQTPGQPPRLLIYRTNYRPTGVPSRFSGAISG
NKAALTITGAQPNDEADYFCTLYKSSANIFGGGTHLTVLGQPKAAPTVNLFPPSSEELGTNKATLVCLISDFYPGAVTVT
WKAGGTTVTQGVETTKPSKQSNNKYAASSYLALSASDWKSSSGFTCQVTHEGTIVEKTVTP…

>IGHG1_C1
…LSVLLMGCVAEEKLVESGGGLVQPGGSLRLSCVGSGFVFSFTYINWVRQAPGKGLEWLAAISTSGGSTYYADSVKGRFT
ISRDNSQNTAYLRMNSLRTEDTALYYCASTLAVDGAMDLWGPGVEVVVSSAPKTAPSVYPLAPCGRDTSGPNVALGCLAS
SYFPEPVTMTWNSGALTSGVHTFPSVLQPSGLYSLSSMVTVPASSLSSKSYT…

>IGK_C1
…LSVLLMGCVAAIVLTQTPLSLSVSPGEPASISCRSSQSLEIYGNNFLSWYQQKPGQSPKLLIYWATNRASGVPDRFSGS
GSGTDFTLTIIRVEAEDAGVYYCQQNKESHMYGFGAGTKLELKRADAKPSVFIFPPSKEQLETQTVSVVCLLNSFFPREV
NVKWKVDGVVQSSGILDSVTEQDSKDSTYSLSSTLSLPTSQYLSHNLYSCEVTHKTLASPLVKSFSRNECEA*

>IGHG1_D8
…LSVLLMGCVAEEKLVESGGGLVQPGGSLRLSCVGSGFTFSNYEINWVRQAPGKGLEWLAAIPNRGGITYYADSVKDRFT
ISKDNSQNTADLQMNSLRTEDTARYYCARSGAGYLYYAMDLWGPGVEVVVSSAPKTAPSVYPLAPCGRDTSGPNVALGCL
ASSYFPEPVTMTWNSGALTSGVHTFPSVLQPSGLYSLSSMVTVPASSLSSKSYTCNVNHPATTTKVDKRVGTKTKPPCPI
CPGCEVAGPSVFIFPPKPKDT…

>IGK_D8
…LSVLLMGCVAAIQLTQSPASLAASLGDTVSITCRASQSINSYLAWYQQQPGKAPKLLIYKASTLQSGVPSRFKGSGSGT
DYTLTISGLQAEDVATYYCLHDNTAPYGFGAGTKLELKRADAKPSVFIFPPSKEQLETQTVSVVCLLNSFFPREVNVKWK
VDGVVQSSGILDSVTEQDSKDSTYSLSSTLSLPTSQYLSHNLYSCEVTHKTLASP…

>IGHG1_F4
…LSVLLMGCVAEEKLVESGGGLGQPGGSLRLSCVGSGFGFSSTYIHWVRQAPGKGLEWLAAISTSVGTTYYYADSVRGRF
TISRDYSQNTAYLQMNSLRTEDTARYYCAREGVSYGDGWDLWGPGVEVVVSSAPKTAPSVYPLAPCGRDTSGPNVALGCL
ASSYFPEPVTMTWNSGALTSGVHTFPSVLQPSGLYSLSSMVTVPASSLSSKSYTC…

>IGK_F4
…LSVLLMGCVAAIQLTQSPASLAASLGDTVSITCRASQSISSYLAWYQQQPGKAPKLLIYYASSLQSGVPSRFKGSGSGT
DFTLTISGLQAEDVATYYCLQHSSAPYGFGAGTKLELKRADAKPSVFIFPPSKEQLETQTVSVVCLLNSFFPREVNVKWK
VDGVVQSSGILDSVTEQDSKDSTYSLSSTLSLPTSQYLSHNLYSCEVTHKTLASPLVKSFSRNEC…

>IGHG1_G9
…LSVLLMGCVAEEKLVESGGELVQPGGSLRLSCVGSGFTFSTTYINWVRQAPGKGLEWLAAVSISSGSTYYADSVKGRFT
ISRDNSQNTAYLQMNSLRTEDTARYYCARGIFGARDIMDLWGPGVEVVVSSAPKTAPSVYPLAPCGRDTSGPNVALGCLA
SSYFPEPVTMTWNSGALTSGVHTFPSVLQPSGLYSLSSMVTVPASSLSSKSYTCNV…

>IGL_G9
…LSVLLMGCVAQTVIQEPAMSVSPGGTVTLTCAFSSGSVTTSNYPSWFQQTPGQPPRLLIYRTNNRPTGVPSRFSGAISG
NKAALTITGAQANDEADYFCTLDKSTSNIFGGGTHLTVLGQPKAAPTVNLFPPSSEELGTNKATLVCLISDFYPGAVTVT
WKAGGTTVTQGVETTKPSKQSNNKYAASSYLALSASDWKSSSGFTCQVTHEGTIVEKTVTPSECA*

Supplementary Figure 5. Analysis of NiV G-specific porcine mAb amino acid sequences. Leader sequences are highlighted yellow, CDR1, CDR2 and CDR3 sequences are highlighted green, cyan and red, respectively, and conserved motifs/residues are highlighted grey.

A

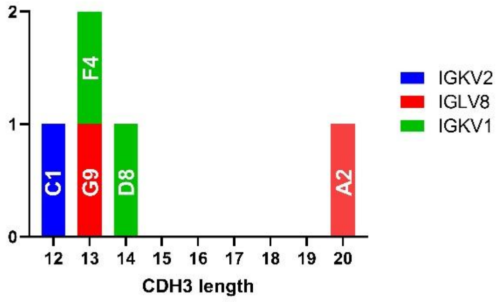


B

| mAb | IGHV distance from germline (bp) | IGLV/IGKV distance from germline (bp) | |
|---|---|---|---|
| | Babraham | CH242 | Sscrofa 11 |
| A2 | 4 | 9 | 14 |
| G9 | 9 | 3 | 6 |
| C1 | 9 | 12 | 14 |
| F4 | 15 | 21 | 23 |
| D8 | 5 | 17 | 19 |

Supplementary Figure 6. (A) CDRH3 lengths and light chain usage of the mAb clones A2, C1, D8, F4, G9. (B) Analysis of somatic hypermutation as measured by distance from known germline sequences: F4 is the furthest from germline both in IGH and IGK. Whereas the IGL clones, A2 and G9, are closest to germline sequences.

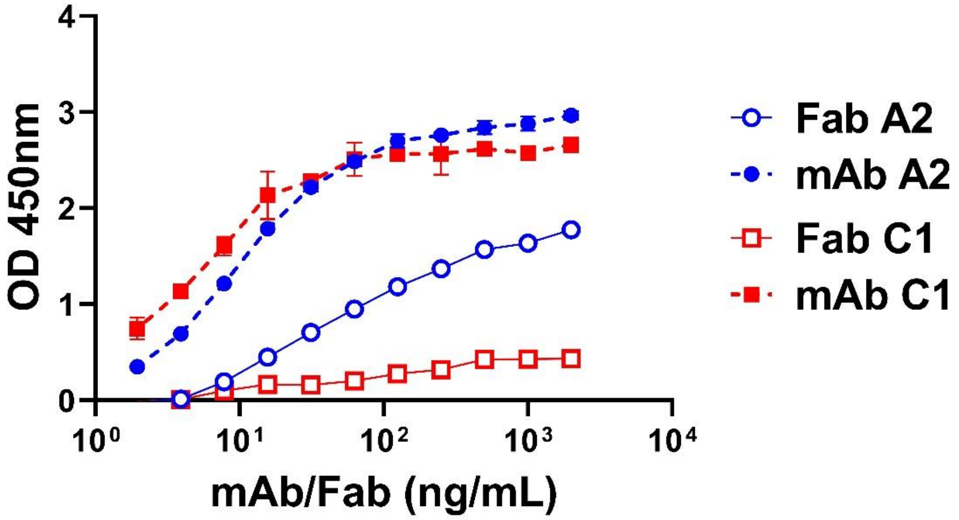


Supplementary Figure 7. Assessment of the binding properties of recombinant NiV G-specific porcine mAbs and Fabs A2 and C1. Recombinant porcine mAbs and Fabs were evaluated for binding to recombinant NiV sG by ELISA. Mean technical duplicate data ±SD from a representative experiment are shown.

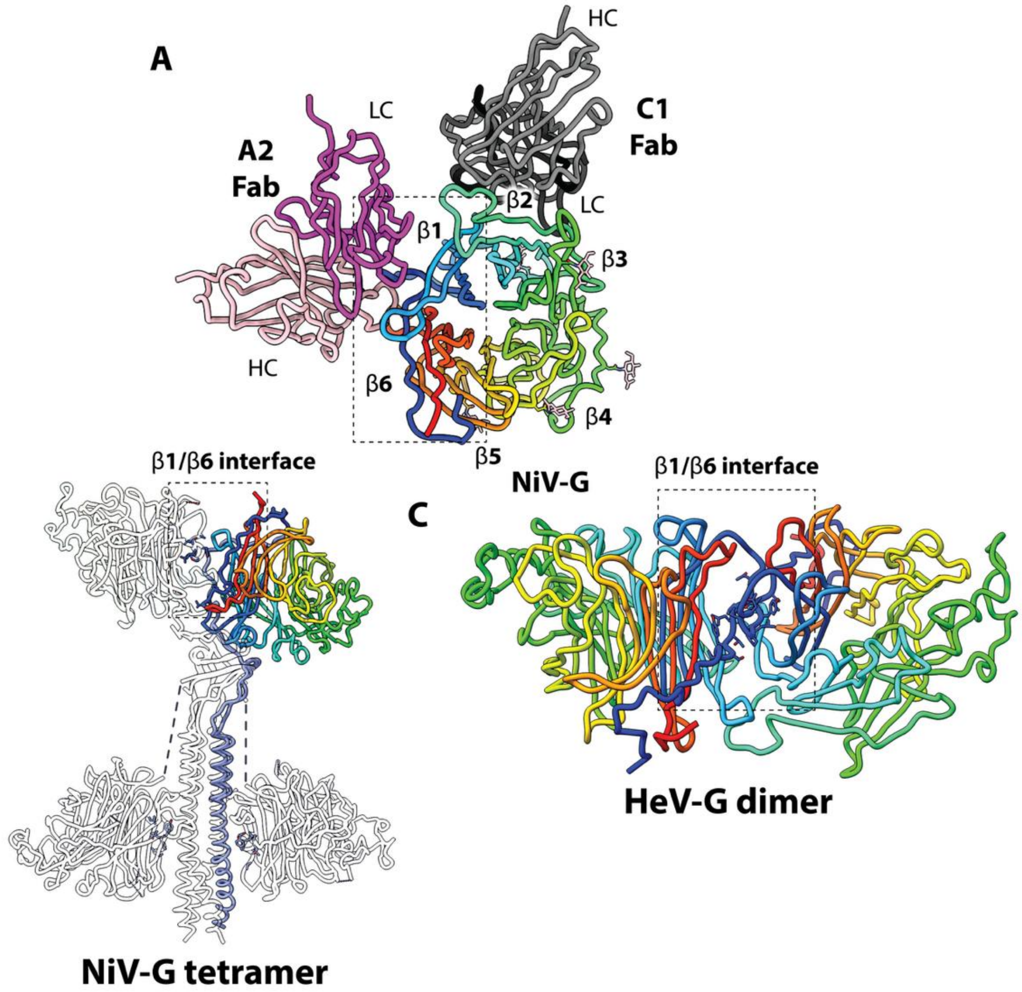


Supplementary Figure 8. The β1/β6 interface responsible for NiV G dimerisation is occluded by mAb A2. The tetrameric cryoEM structure of NiV G (PDB: 7TXZ and 7TY0; [14]) and dimeric crystal structure of HeV G (PDB: 2X9M; [17]) are shown in cartoon loop orientation. For (B), individual protomers are transparent with sharp outline except one protomer which has purple coloured stalk and rainbow coloured RBD (residues 183-602).

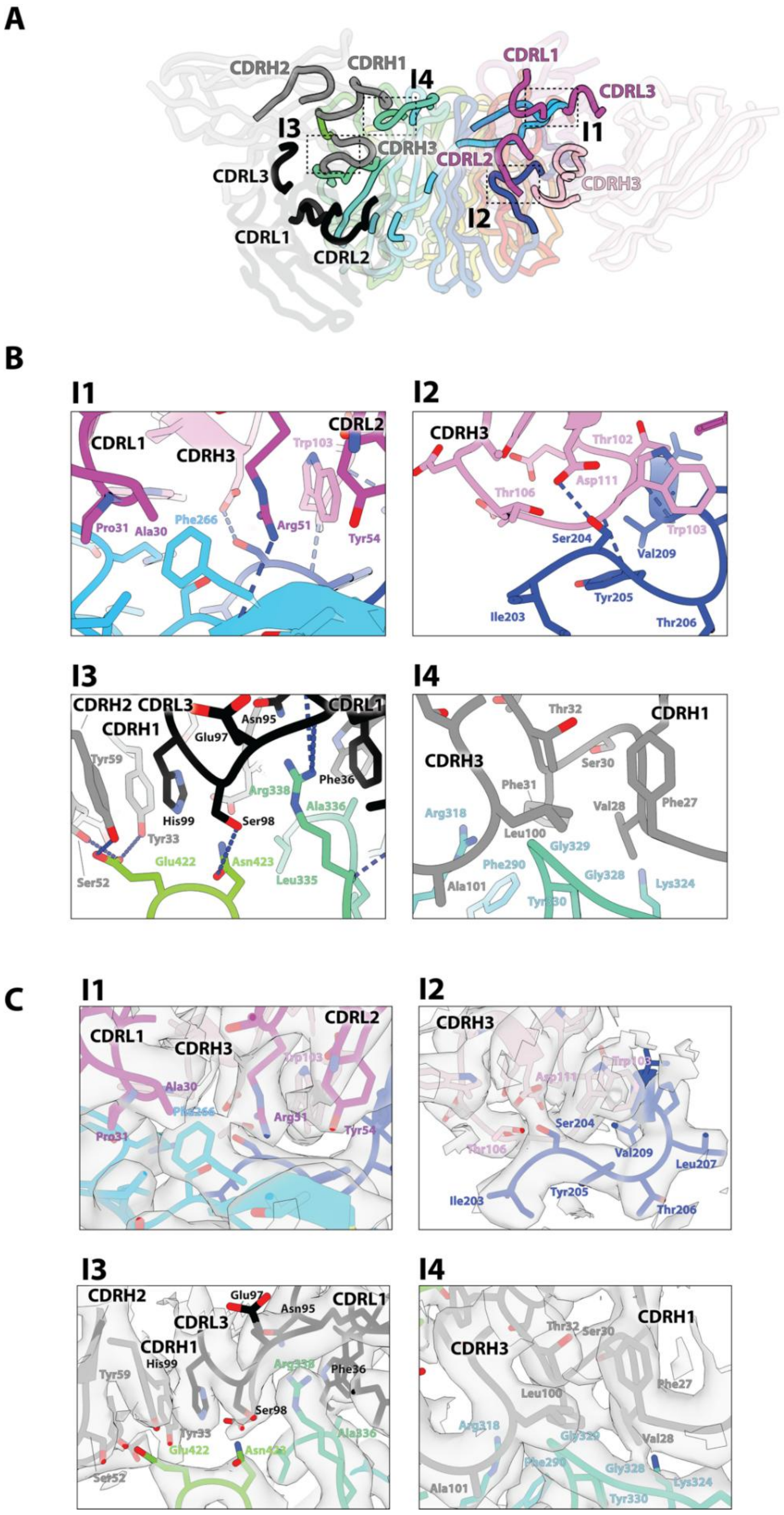


Supplementary Figure 9. Interaction between Fabs A2 and C1 with NiV G. (A) View of the junction between β1 and β2. CDR loops are coloured grey (C1 heavy), black (C1 light), magenta (A2 light), and pink (A2 heavy). Antibody-NiV G interface regions (I1–I4) are highlighted in dotted frames. (B) Close-up views of the interaction sites (I1-I4) with the same colouring as (A). Relevant amino acid sidechains are presented as sticks. (C) The same close-up views as (B) with cryoEM density present for relevant residues. The map is contoured at level = 0.40.

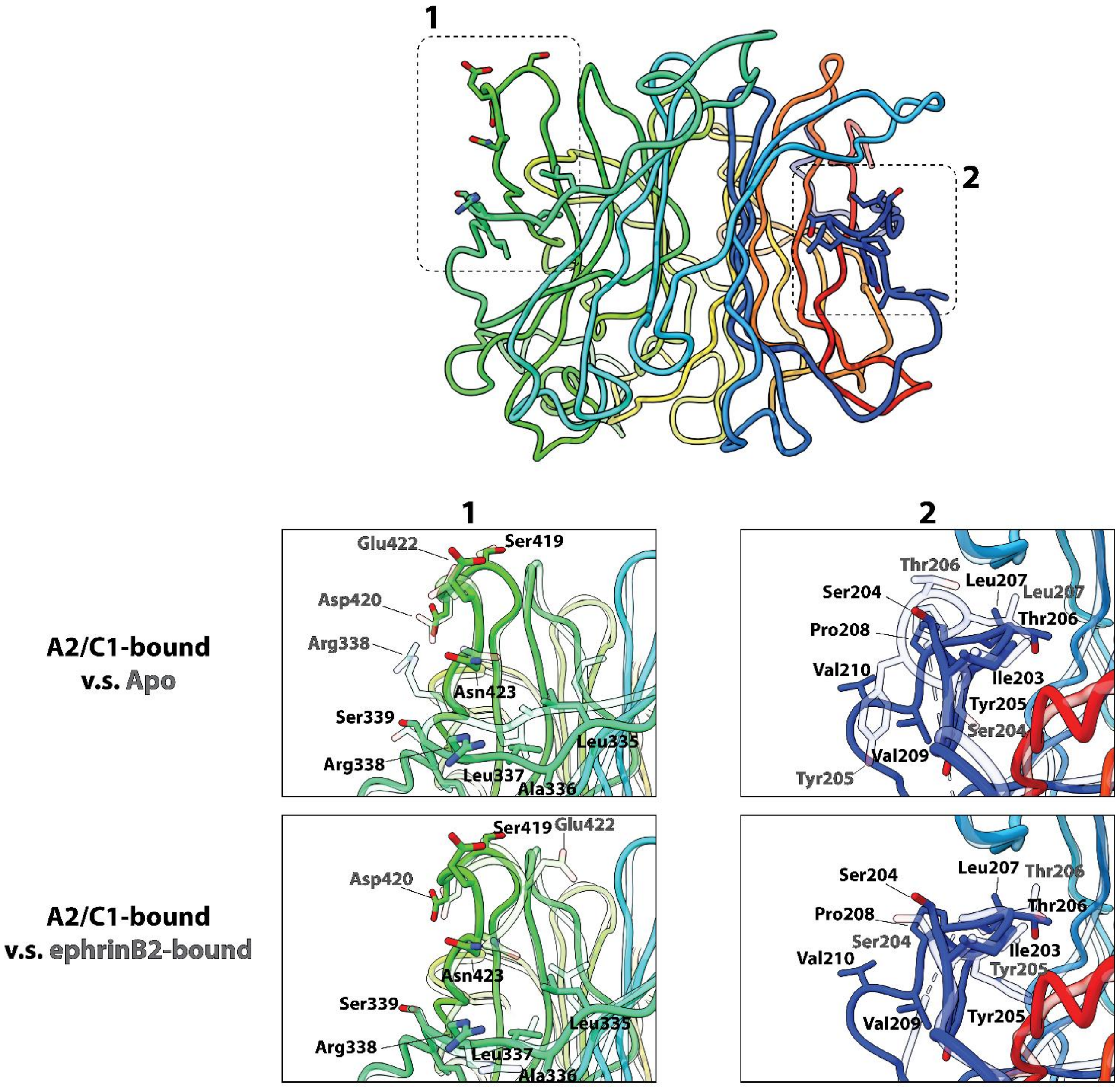


Supplementary Figure 10. Fabs A2 and C1 induce a ‘receptor-bound’ like conformation. Apo and ephrinB2-bound structures are taken from PDB: 2VWD [18] and 2VSM [19], respectively. The overlayed apo or ephrinB2-bound structures are presented as transparent cartoons with sharp black outline.

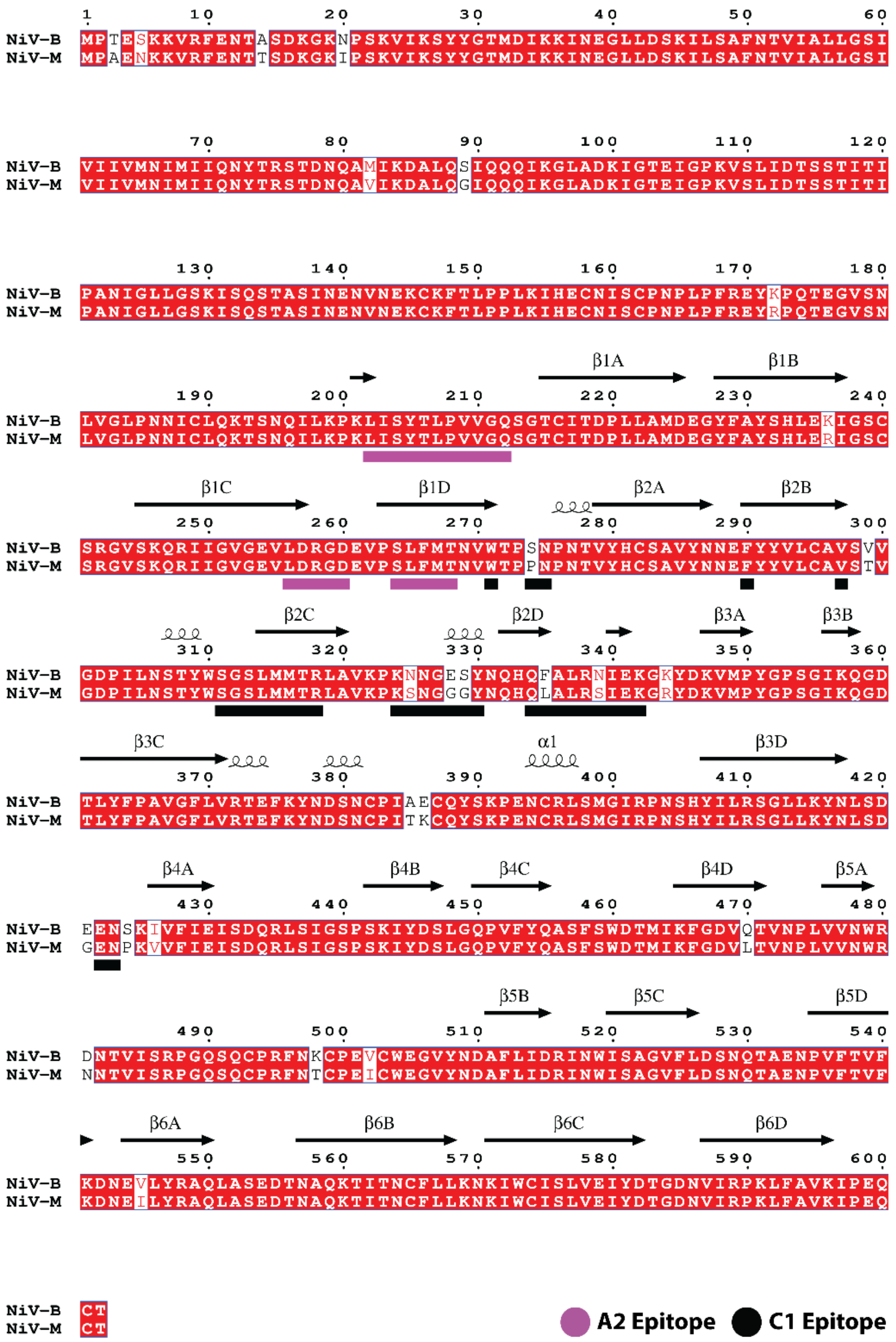


Supplementary Figure 11. Sequence alignment between NiV-M and NiV-B G with mAb A2 and mAb C1 epitopes highlighted. Sequences were retrieved from Uniprot: Q4VCP5 (NiV-B) and Q9IH62 (NiV-M), aligned with Clustal-O webserver [20], and visualised with ESPript webserver [21]. A2 and C1 interacting residues on NiV-M are highlighted with purple and black stripes underneath. Conserved residues between NiV-M and NiV-B are highlighted in red. Secondary structures including blade nomenclature for the β-propeller are annotated above the sequences block.

Supplementary Table 2. Survey of structurally characterised anti-NiV G-specific mAbs.

| ***mAb*** | ***Epitope*** | ***Number of mAbs from Epitope binning*** | ***Origin*** | ***Immunogen used to elicit mAb*** | ***Refs*** |
|---|---|---|---|---|---|
| *m102.3* | RBS | 8 | Human Fab phage-display library | None | [4-7] |
| *HENV-26* | RBS | 1 | Human | HeV sG | [8] |
| *1E5* | RBS | 4 | Macaque | rAd5-NiV-B G | [9] |
| *14F8* | RBS | 1 | Murine | NiV-B sG | [10] |
| *LN1F9* | RBS | 7 | Murine | NiV sG-apoferritin | [11,12] |
| *HENV-32* | DI | 4 | Human | HeV-sG | [8] |
| *A2* | DI | 1 | Porcine | NiV sG mRNA | This publication |
| *nAH1.3* | NS | 1 | Murine | Virus | [13,14] |
| *LN3D3* | NS | 16 | Murine | NiV sG-apoferritin | [11,12] |
| *S2B10* | β1 | 1 | Murine | NiV sG-apoferritin | [11,12] |
| *C1* | β2/β3 | 1 | Porcine | NiV sG mRNA | This publication |
| *S1E2* | β3 | 3 | Murine | NiV sG-apoferritin | [11,12] |

RBS = receptor-binding site, DI = dimerisation interface, NS = near-stalk region
